# Dietary methionine restriction targets one carbon metabolism in humans and produces broad therapeutic responses in cancer

**DOI:** 10.1101/627364

**Authors:** Xia Gao, Sydney M. Sanderson, Ziwei Dai, Michael A. Reid, Daniel E. Cooper, Min Lu, John P. Richie, Amy Ciccarella, Ana Calcagnotto, Peter G. Mikhael, Samantha J. Mentch, Juan Liu, Gene Ables, David G. Kirsch, David S. Hsu, Sailendra N. Nichenametla, Jason W. Locasale

**Affiliations:** Department of Pharmacology and Cancer Biology, Duke University School of Medicine, Durham, NC 27710, USA; Department of Radiation Oncology, Duke University Medical Center, Durham, NC 27710, USA; Center for Genomics and Computational Biology, Duke University, Durham, NC 27710, USA; Department of Medical Oncology, Duke University Medical Center, Durham, NC 27710, USA; Penn State University College of Medicine, Department of Public Health Sciences, Hershey, PA 17033, USA; Penn State University Clinical Research Center, State College, PA 16802, USA; Orentreich Foundation for the Advancement of Science, Cold Spring, NY 10516, USA

## Abstract

Nutrition exerts profound effects on health and dietary interventions are commonly used to treat diseases of metabolic etiology. Although cancer has a substantial metabolic component, the principles that define whether nutrition may be used to influence tumour outcome are unclear. Nevertheless, it is established that targeting metabolic pathways with pharmacological agents or radiation can sometimes lead to controlled therapeutic outcomes. In contrast, whether specific dietary interventions could influence the metabolic pathways that are targeted in standard cancer therapies is not known. We now show that dietary restriction of methionine (MR), an essential amino acid, and the reduction of which has aging and obesogenic properties, influences cancer outcome through controlled and reproducible changes to one carbon metabolism. This pathway metabolizes methionine and further is the target of a host of cancer interventions involving chemotherapy and radiation. MR produced therapeutic responses in chemoresistant RAS-driven colorectal cancer patient derived xenografts and autochthonous *KRAS*^*G12D*+/−^;*TP53*^−/−^ -driven soft tissue sarcomas resistant to radiation. Metabolomics revealed the therapeutic mechanisms to occur through tumor cell autonomous effects on the flux through one carbon metabolism that impacted redox and nucleotide metabolism, thus interacting with the antimetabolite or radiation intervention. Finally, in a controlled and tolerated feeding study in humans, MR resulted in similar effects on systemic metabolism as obtained in responsive mice. These findings provide evidence that a targeted dietary manipulation can affect specific tumor cell metabolism to mediate broad aspects of cancer outcome.

## Main

The dietary intake of nutrients, and ultimately the levels of circulating metabolites that provide nutrients to tumors are determined by a manifestation of complex physiological processes occurring in liver, gut, pancreas, brain, and other tissues. While it is known that changes in the growth media that cancer cells encounter in culture can have dramatic effects on cell metabolism and cell fate ^1–3^, the extent to which dietary nutrient availability, which is the corresponding situation *in vivo*, can alter metabolic pathways in tumors and affect therapeutic outcomes in cancer is largely unknown. Recent studies ^4–6^ have shown that the restriction of amino acids, serine and glycine, can modulate cancer outcome in xenotransplanted and autochthonous tumor models. The availability of histidine and asparagine appears to mediate the response of cancer cells to methotrexate ^7^ and the progression of breast cancer metastasis ^8^, respectively. Whether such interventions broadly affect metabolism or have targeted effects on specific pathways related to these nutrients is unknown. One intriguing possibility for a specific dietary intervention in cancer is the restriction of methionine, an essential amino acid in one carbon metabolism. Methionine is the most variable metabolite found in human plasma ^9^, has a myriad of functions as a result of its location in one carbon metabolism ^10^, and dietary MR is known to extend lifespan ^11–15^, and improve metabolic health^16–19^. Of particular interest, one carbon metabolism, through its essential role in redox and nucleotide metabolism, is the target of frontline cancer chemotherapies such as 5-fluorouracil, antifolates (e.g. methotrexate and pemetrexed), and radiation therapy ^20–22^. Thus some cancer cell lines exhibit differing extents of methionine auxotrophy ^23,24^ and depleting or restricting methionine from the diet may possibly have anti-cancer effects in mice ^25–28,29,30^. Therefore, we reasoned that MR could have broad anti-cancer properties by targeting a focused area of metabolism that would interact with the response to other therapies that also affect one carbon metabolism.

### Dietary MR rapidly and specifically alters plasma methionine and sulfur metabolism

We and others have previously reported that MR alters metabolism in mouse liver and plasma after a long-term intervention ^9^, but its impact on metabolism on acute time scales that could affect metabolic health and tumor growth in treatment settings is less explored. To study the dynamics of dietary MR on systemic metabolism, we switched the diet of C57BL/6J male mice from a chow diet to a control diet (0.86% methionine w/w) or a MR diet (0.12% methionine w/w) as previously defined ^9^ for three weeks, and obtained metabolite profiles of plasma at days 0, 1, 2, 4, 7, 10, 14, 17, and 21 (Figure 1A) using a comparative metabolomics approach ^9,20^. With a singular value decomposition ^31^, the time course was decomposed into a series of rank-ordered modes that define the relative contributions of sets of metabolites that correspond to different sources of metabolic dynamics (Figure 1B and S1A). While Mode 1 reflected an overall change in metabolism due to switching diets at time zero, Modes 2 and 3 predominantly contained metabolites related to methionine and sulfur metabolism (Figure 1B-C). Hierarchical clustering confirmed that a set of methionine-related metabolites was most rapidly suppressed with other compensatory pathways changing at later times (Figure 1D). By the 3-week end point, MR showed predominant alterations in cysteine and methionine metabolism, followed by taurine and hypotaurine metabolism, and tryptophan metabolism (Figures 1E-F, S1B). Methionine was the only amino acid in plasma significantly reduced by MR (Figure S1 and Table S1). Dramatically, metabolites in methionine metabolism such as methionine, methionine sulfoxide, and 2-keto-4-methylthiobutyrate were reduced over 50% within one day of MR and such reduction remained at similar levels throughout the dietary intervention (Figure 1G). Hypotaurine involved in the transsulfuration pathway was also decreased after two days of MR (Figure 1G). Redox balance at the 3-week end point was largely maintained (Figure S1B). Together, these findings show that dietary MR rapidly and reproducibly alters circulating metabolism in mice with sustained effects concentrated primarily on metabolism related to methionine.

**Figure 1.**
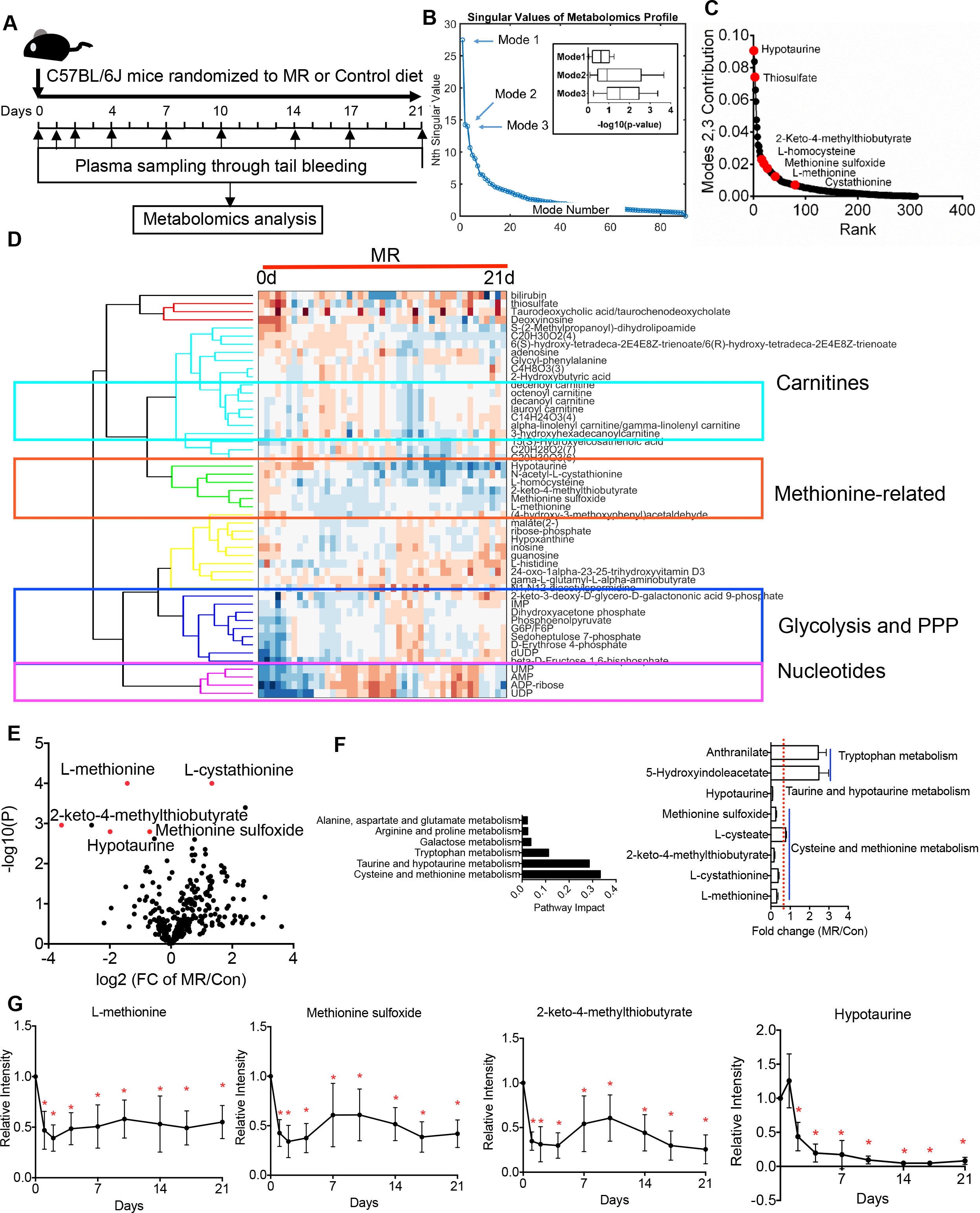
Dietary MR rapidly and specifically alters methionine and sulfur metabolism but maintains overall metabolism. A, Schematic of experimental design. 12-week-old male C57BL/6J mice were subjected to either the control or the MR diet *ad libitum* for 3 weeks (n=5/group). Blood was sampled through tail bleeding at day 0, 1, 2, 4, 7, 10, 14, 17, 21 post-dietary treatments. B, 90 sets of metabolic profiles from mice in A (presented in a heatmap) were computed for singular values via singular value decomposition (SVD). Insertion: T-test p-values assessing difference between control and MR in the first three modes. C, Contribution of mode 2 and mode 3 in B ranked from highest to lowest across all measured metabolites. D, Heatmap of metabolites in mode 2 and mode 3 of B from mice on the MR diet. E, Volcano plot of metabolites in plasma collected at the end point. F, Left: Pathway analysis of significantly changed (*p<0.05, two-tailed Student’s t-test) plasma metabolites by 21-day MR diet versus that from mice on the control diet. Right: Fold change (FC) of altered metabolites in cysteine and methionine metabolism, taurine and hypotaurine metabolism and tryptophan metabolism. G, Relative intensity of selected metabolites in cysteine and methionine metabolism over time. Data are represented as mean ± SD. *p<0.05.

### Dietary MR inhibits tumor growth and affects methionine metabolism in colorectal patient-derived xenograft (PDX) models

Given that MR rapidly and specifically affects systemic methionine metabolism, we sought to evaluate its impact in a series of cancer settings related to one carbon metabolism. PDX models recapitulate genomic features of their original tumors in an *in vivo* environment ^32,33^. Thus, to test whether MR may have an impact on tumor growth, we first considered two RAS-driven colorectal cancer (CRC) PDX models (Figure S2A) ^34,35^: CRC119 derived from a patient bearing a KRAS^G12A^ mutation and CRC240 carrying a NRAS^Q61K^ mutation. We set up two regimens (Figure 2A): in experiment 1, mice were subjected to the control diet or the MR diet when the tumor was palpable (treatment); and in experiment 2, mice were subjected to the above dietary treatment two weeks prior to inoculation (prevention). Tumor growth in male and female mice in both CRC PDX models responded similarly to dietary MR: MR alone inhibited the growth of established tumors in CRC 119 PDX model (p=5.71e-12 at the end point, two-tailed Student’s t-test), and showed a trending effect in CRC240 model (p=0.054 at the end point, two-tailed Student’s t-test) (Figure 2B), and pre-dietary treatment led to a stronger effect in each case (Figure 2C). Interestingly, mice on MR exhibited similar or higher amounts of food intake than for the mice fed the control diet (Figure S2B), implying the inhibitory effect of dietary MR on tumor growth is not due to caloric restriction. Together, these data suggest a possible therapeutic potential and a preventive function of dietary MR in RAS-driven CRCs.

**Figure 2.**
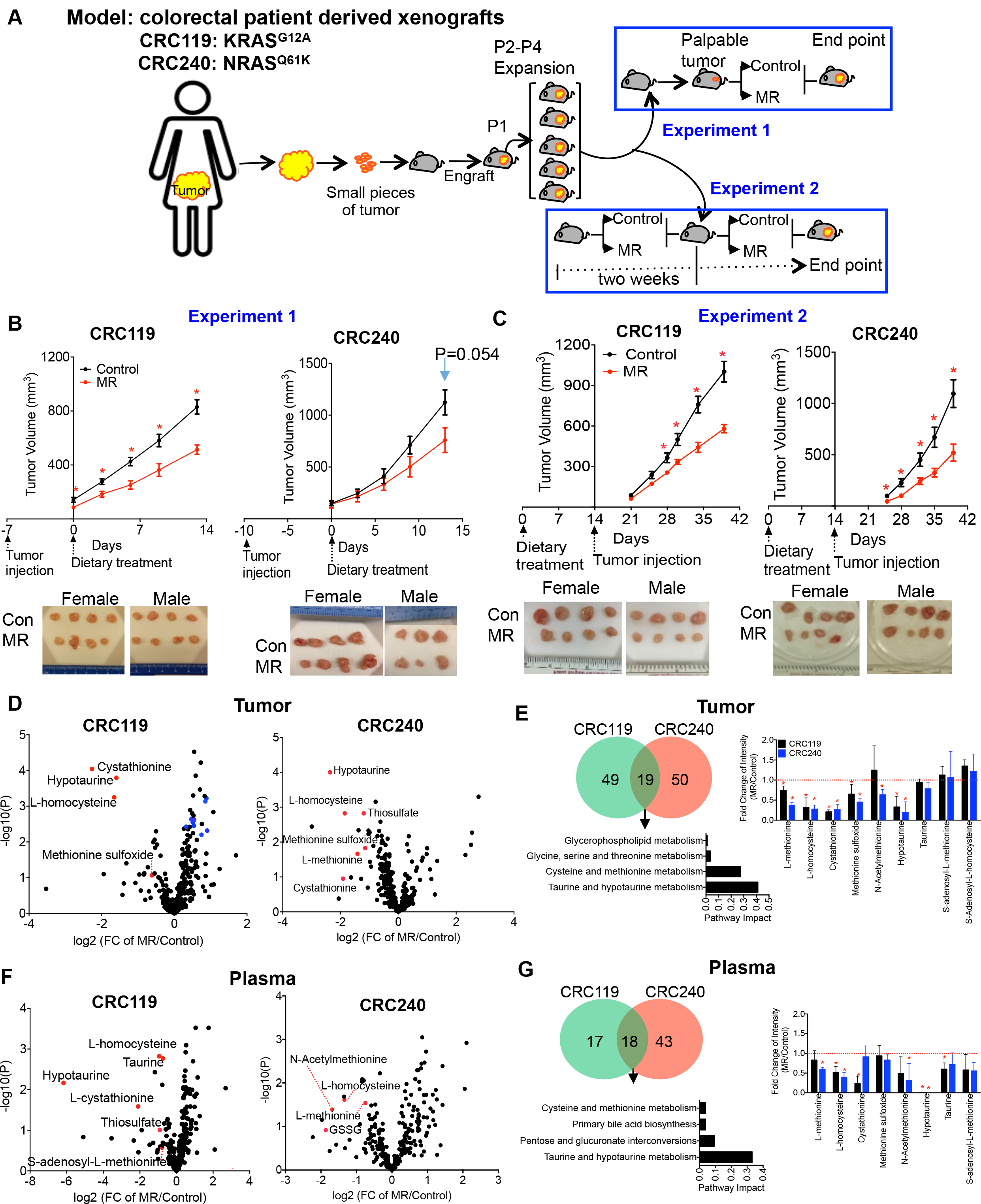
Dietary MR inhibits tumor growth and affects methionine metabolism in colorectal patient-derived xenograft (PDX) models. A, Schematic of experimental design. Two colorectal PDXs were used. Patient-derived tumors were passed through 8-10-week-old JAX NOD.CB17-PrkdcSCID-J mice for 2-4 generations. In experiment 1, mice were injected with tumor first and then randomly subjected to the control or the MR diet when tumor was palpable (n=7/group, 4 females + 3 males). In experiment 2, mice were randomly subjected to the control or the MR diet 2 weeks prior to tumor inoculation until tumor reached the end point (n=8/group, 4 females + 4 males). B, Tumor growth curve and images of tumors at the end point from experiment 1. Data are represented as mean ± SEM. *p<0.05 by two-tailed Student’s t-test. C, Tumor growth curve and images of tumors at the end point from experiment 2. Data are represented as mean ± SEM. *p<0.05, two-tailed Student’s t-test. D, Volcano plots of metabolites in tumors from C. Blue dots indicate acylcarnitines. E, Left: Venn diagram of significantly changed (*p<0.05, two-tailed Student’s t-test) metabolites in tumors by MR from C and pathway analysis (false discovery rate < 0.5) of the commonly changed metabolites in both models; right: MR-induced fold change (FC) of tumor metabolites in cysteine and methionine metabolism, and taurine and hypotaurine metabolism. Data are represented as mean ± SD. *p<0.05. F, Volcano plots of metabolites in plasma from C. G, Left: Venn diagram of significantly changed (*p<0.05, two-tailed Student’s t-test) metabolites in plasma by MR from C and pathway analysis of the commonly changed metabolites in both models; right: MR-induced fold change (FC) of tumor metabolites in cysteine and methionine metabolism, and taurine and hypotaurine metabolism. Data are represented as mean ± SD. *p<0.05.

To gain insights into metabolism in these settings, we profiled metabolites in tumor, plasma and liver from mice having undergone pre-dietary treatment. As seen in non-tumor bearing mice, dietary MR prominently altered methionine and sulfur-related metabolism across all examined tissue types (Figures 2D-G and S2C-D). In tumors, MR decreased the levels of methionine (suggesting a possibility of decreased protein synthesis) and related metabolites such as homocysteine, cystathionine, methionine sulfoxide, and hypotaurine in both CRC119 and CRC240, but not S-adenosylmethionine (SAM) (Figures 2E and S2E, and Table S1). A similar pattern was also observed in plasma and liver (Figures 2F and S2D-E). Unaltered levels of SAM and S-adenosylhomocysteine (SAH) were observed in the tumors (Figures 2E) suggesting that methylation reactions may not be compromised. Together, dietary MR inhibits tumor growth and consistently alters methionine and sulfur metabolism in tumor, plasma, and liver in RAS-driven CRC PDX models.

### MR induces tumor cell intrinsic metabolic alterations

Dietary MR altered methionine metabolism in tumor, liver, and plasma, but it is not clear whether the effect on tumor growth is systemic, cell autonomous, or both. To address this question, we conducted an integrated analysis of global changes in the metabolic network across tumor, plasma and liver within each PDX model. The fold changes (FC) of metabolites measured across all of metabolism exhibited strong correlations between each pair corresponding to tumor, plasma and liver (Figures 3A-B) and showed the highest correlation between tumor and plasma, with Spearman rho of 0.51 (p=1.37e-15) and 0.29 (p=1.78e-5) for CRC119 and CRC240 respectively (Figures 3A-B). To investigate the relationship between the FC of metabolites among tumor, plasma and liver, we employed a multidimensional scaling analysis (methods). In both models, the FC of metabolites in tumor showed a higher similarity with those in plasma than with those in liver (Figure 3C). Liver was the most affected tissue in comparison to tumor and plasma in both PDX models (Figure 3D-E). To further determine the specificity of MR on metabolism among tumor, plasma and liver, we classified metabolites in all examined tissue types into either methionine-related or methionine-unrelated. We defined metabolites as methionine-related if the metabolite was metabolized from or to methionine within 4 reaction steps in the human metabolic network ^36^, and methionine-unrelated meaning that the compound requires more than 4 reactions to be converted from or to methionine (Figure 3F). In plasma and tumor, a higher proportion of the altered metabolites was methionine-related in both models (Figure 3G-H). In contrast, MR-altered metabolites in liver were nearly equally distributed between methionine-related and -unrelated metabolite groups (Figure 3G). These data suggests that cell autonomous effects of MR on tumor metabolism that originate from suppressed plasma methionine levels. Consistently in patient-derived primary tumor cells, MR inhibited proliferation of cells from both PDX models (Figure S3A) and most significantly altered metabolites in cysteine and methionine metabolism (Figure S3B-C) as in the PDX tumors. In both cell lines, MR led to reduction in methionine, 5’-methylthioadenosine, methionine sulfoxide, hypotaurine, taurine and SAM (Figure S3C-D). Thus, the inhibition of tumor growth is at least partially attributed to a cell autonomous effect of MR on tumor.

**Figure 3.**
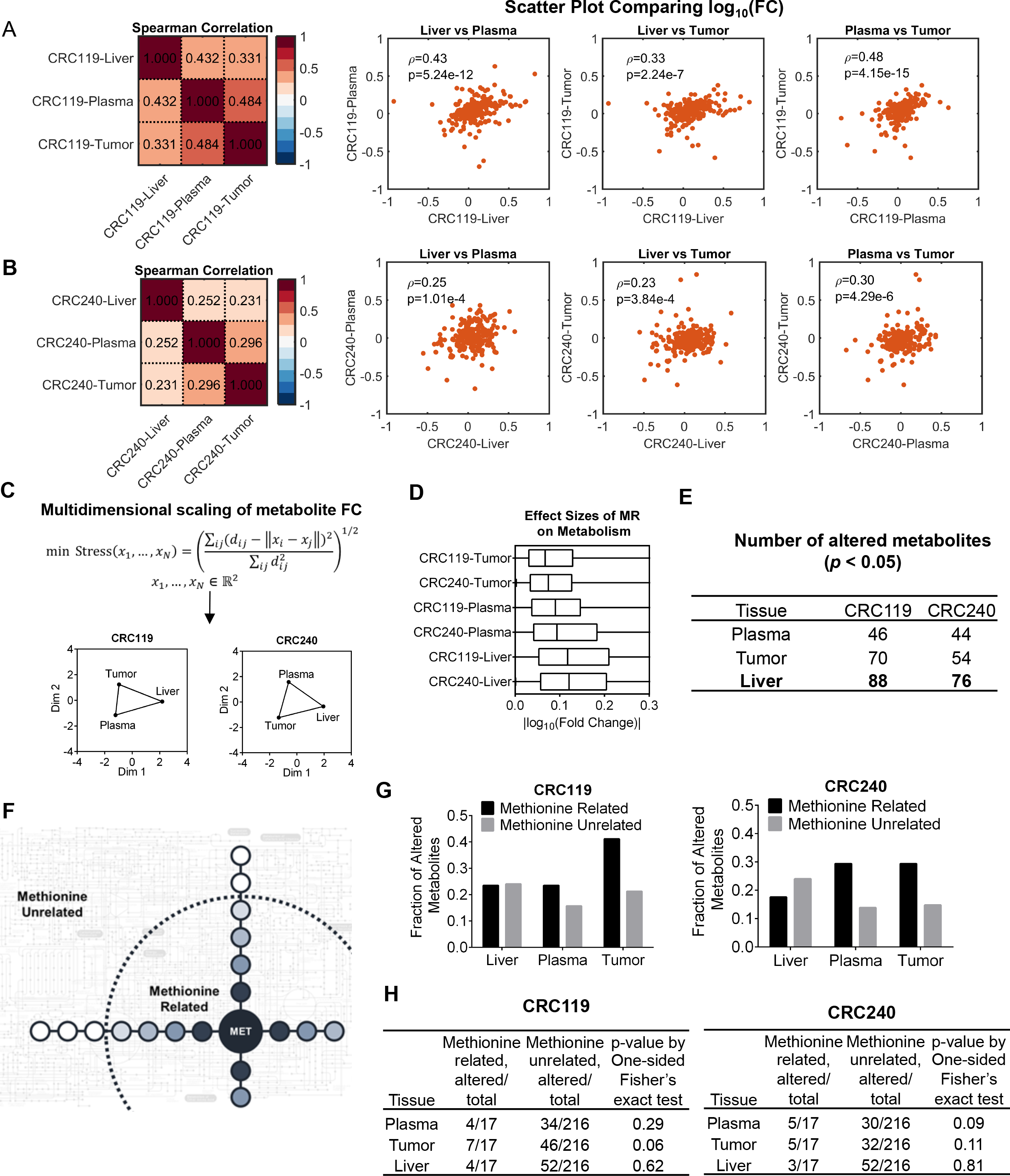
MR leads to specific cell intrinsic metabolic alterations in tumor. A, Spearman’s rank correlation coefficients of MR-induced fold changes (FC) of metabolites in tumor, plasma and liver from CRC119 model in Figure 2C. B, Spearman’s rank correlation coefficients of MR-induced FC of metabolites in tumor, plasma and liver from CRC240 model in Figure 2C. C, Multidimensional scaling of metabolite FC in response to MR in CRC119 and CRC240 models. D. Effect of MR on metabolism in tumor, plasma and liver from CRC119 and CRC240 models in Figure 2C evaluated by the log_10_ (Fold change). E, Numbers of metabolites significantly altered (*p<0.05, two-tailed Student’s t-test) by MR in plasma, tumor and liver in CRC119 and CRC240 models F, Schematic defining methioninerelated (metabolized from or to methionine within 4 reaction steps) metabolites and otherwise methionine non-related metabolites. G, Fraction of significantly altered metabolites for methionine-related and methionine-unrelated metabolites in tumor, liver and plasma from CRC119 and CRC240 models in Figure 2C. H, Numbers of total and significantly altered metabolites for methionine-related and methionine-unrelated metabolites, and the p values analyzed by one-sided Fisher’s exact test in tumor, liver and plasma from CRC119 and CRC240 models in Figure 3G.

### Dietary MR sensitizes CRC PDX models to 5-Fluorouracil (5-FU)

5-FU targets thymidylate synthase, an enzyme related to methionine and one carbon metabolism ^20^, and is a frontline chemotherapy for colorectal cancer with therapeutic strategies in CRC achieving modest (~60-65%) responses ^37,38^. We therefore tested whether MR could synergize with 5-FU (Figure 4A). Considering the reproducible inhibition of MR on tumor growth and the common metabolic effects of MR, we further explored this possibility in the CRC119 model. To minimize the toxicity, we delivered an established low dose of 5-FU ^39^ to CRC119 tumor-bearing mice, and such dose showed no effect on tumor growth in mice fed the control diet (Figure 4B). However, MR synergized with 5-FU, leading to a marked effect on tumor growth in both male and female mice (Figure 4B). The combination of MR and 5-FU led to a broad effect on metabolic pathways in tumor, plasma and liver (Figure 4C and S4A-D), with the most prominent changes centered on nucleotide metabolism in tumors related to both the mechanism of action of 5-FU and MR (Figure 4C-D). 5-FU treatment increased nucleoside levels including guanosine, cytidine, uridine and thymidine, but reduced nucleotides with nucleobase levels largely maintained (Figure 4D). Notably, these patterns are consistent with inhibition of thymidylate synthase and a global disruption to de novo nucleotide synthesis, thus confirming that with MR, 5-FU is able to engage targets effectively likely due to limitations in one carbon metabolism created by lower activity in the methionine cycle. In addition, the combination of 5-FU and MR largely disturbed cellular redox balance in the tumor, reflected by a decrease of glutathione (GSH), NADH, citrate, a decreased ratio of GSH/GSSG (an oxidized form of GSH), a reduced α-ketoglutarate/citrate ratio, and an increased ratio of NADH/NAD^+^ (Figure 4E). Interestingly in the liver, the combined therapy led to a reduction of NAD+, NADH, but not GSH or a decreased ratio of GSH/GSSG. Contrary to the reduced α-ketoglutarate/citrate ratio and increased ratio of NADH/NAD in the tumor, we saw an increased α-ketoglutarate/citrate ratio and no change in the ratio of NADH/NAD in the liver (Figure S4F). Meanwhile in the plasma, there was no significant change in glutathione, oxidized glutathione, citrate, α-ketoglutarate, or the ratio of GSH/GSSG and α-ketoglutarate/citrate (Figure S4F). Thus, the impact of MR on redox balance is tumor specific. A comparative analysis at the global level across the metabolic network revealed that the FC of metabolites between plasma and liver were highly correlated (Spearman’s rho = 0.38, p = 6.7e-11) but that tumor profile exhibited little correlation with systemic metabolism as measured from liver (Spearman’s rho = 0.14, p = 0.02) and circulation (Spearman’s rho = 0.14, p = 0.03) (Figure S4E). Together, dietary MR synergizes with 5-FU, inhibiting tumor growth in CRC PDX models and disrupts nucleotide metabolism and redox balance that underlie the increased efficacy of the antimetabolite chemotherapy.

**Figure 4.**
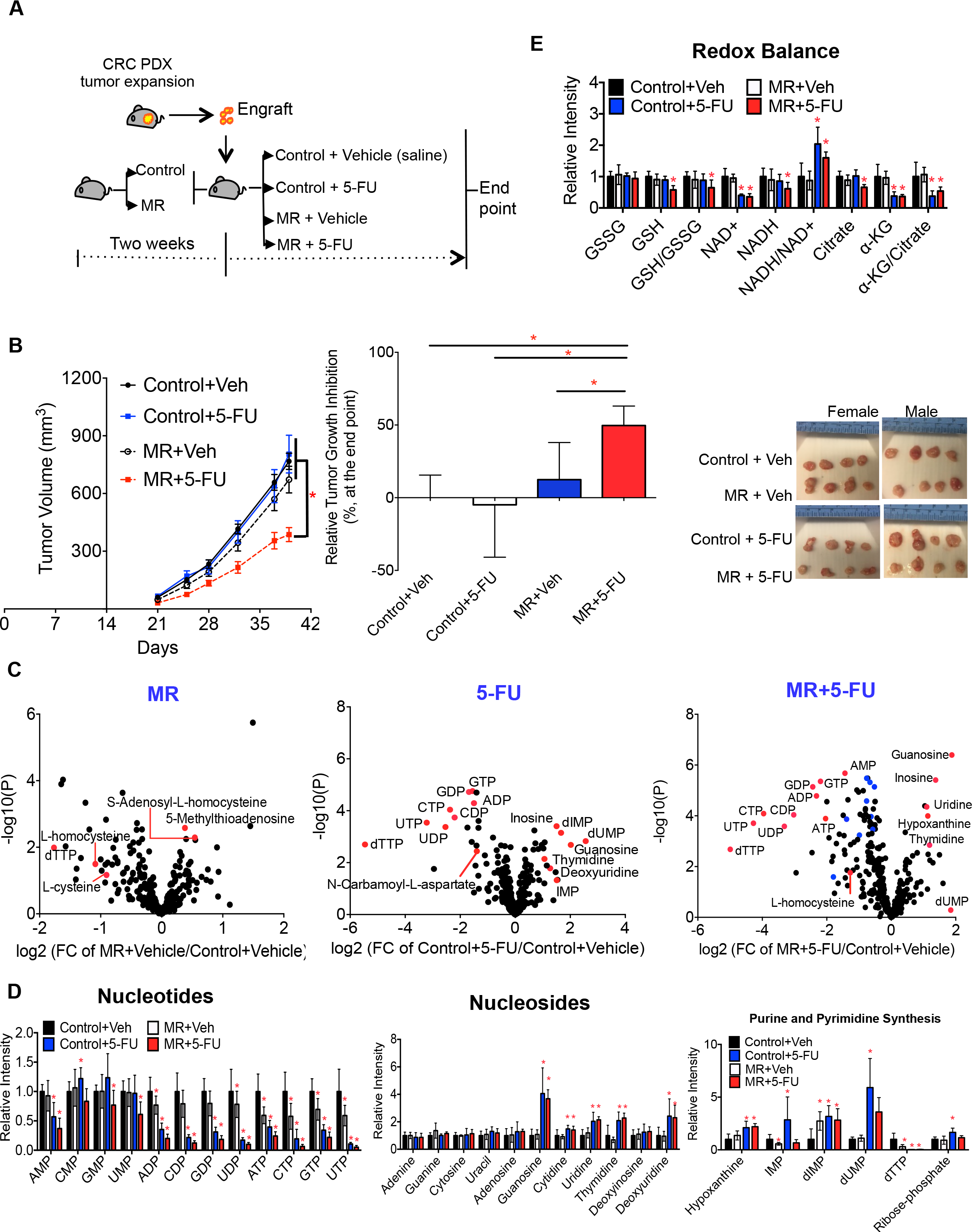
Dietary MR sensitizes CRC PDX models to chemotherapy 5-Fluorouracil (5-FU) A, Schematic of experimental design. PDX model CRC119 was used. Mice were randomly subjected to the control or the MR diet prior to tumor inoculation. 5-FU (12.5 mg/kg × 3 times/week) or vehicle (saline) were intraperitoneally delivered when tumor was palpable. Study ended when tumor reached its end point. (n=8/group, 4 females + 4 males) B, Tumor growth curve, tumor inhibition index and tumor image at the end point. Data are represented as mean ± SEM. *p<0.05. C, Volcano plots of metabolites in tumor altered by MR, 5-FU, or by the combination of dietary MR and 5-FU. FC, fold change. Blue dots indicate acylcarnitines. D, Relative intensity of nucleotides and nucleosides. Data are mean ± SD. *p<0.05 compared to the control group by two-tailed Student’s t-test. E, Relative intensity of metabolites related to redox balance and the ratio of GSH/GSSG, NADH/NAD+ and α–KG/Citrate in tumors. Data are mean ± SD. *p<0.05 compared to the control group by two-tailed Student’s t-test. GSH, glutathione; GSSG, the oxidized form of glutathione; α-KG, α-ketoglutarate.

### MR disrupts cell proliferation by targeting one carbon metabolism

To further study the mechanisms of MR-mediated inhibition of tumor cell proliferation, we first considered “rescue” experiments in cell culture using primary CRC119 cells and HCT116 colorectal cancer cells that are sensitive to MR and 5-FU ^20^. We evaluated the effects of supplementation of a suite of nutrients related to methionine metabolism and the observed differences in metabolite profiles found in the mouse models, including one-carbon donors, choline and formate; sulfur-donor homocysteine with or without cofactor Vitamin B12; nucleosides, and antioxidant N-acetyl cysteine (NAC) on their ability to rescue MR-caused defects in cell proliferation (Figure 5A). We employed a MR condition, 10 μM of methionine that is one tenth of the amount in RPMI media, to approximately model the differences in dietary methionine we consider that was sufficient to inhibit cell proliferation (Figure 5B). While choline and homocysteine alone showed no effects, addition of formate, nucleosides, and NAC all partially alleviated the inhibition of cell proliferation in CRC119 cells (Figure 5B). Addition of homocysteine and B12 also showed a trend towards partial rescue in CRC119 cells. Of great interest, a combination of homocysteine, B12, nucleosides, and NAC fully rescued the MR-mediated inhibition of cell proliferation. These observations were largely replicated in HCT116 colorectal cancer cells. These data are in line with our observation *in vivo* and indicate that disruptions to nucleotide metabolism and cellular redox balance underlie the effects of MR on cancer cell proliferation.

**Figure 5.**
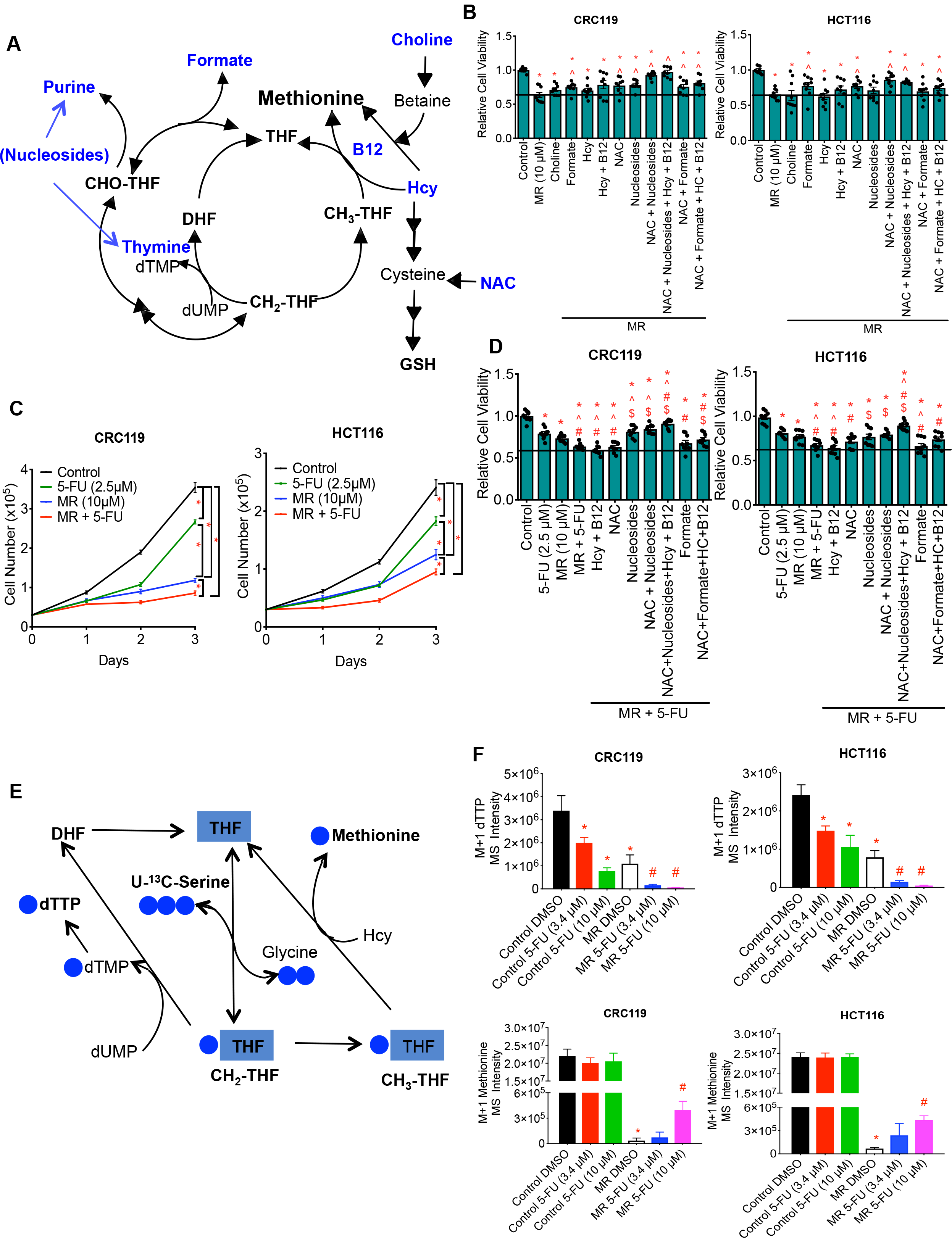
MR-mediated inhibition of cell growth is largely due to interruptions to nucleosides production and redox balance. A, Schematic of rationale for rescue experiments: metabolites used for rescue are in blue. B, Rescue effect of choline, formate, sulfur-donor homocysteine (Hcy), Hcy+B12, nucleosides, and antioxidant N-acetyl cysteine (NAC) alone or in combination on MR-mediated inhibition of cell proliferation. 5-7.5 ×10^3^ cells were plated in 96-well plate. Relative cell numbers were evaluated by MTT assay after 48 h of treatment. Data are mean ± SEM. * p<0.05 vs control and ^ p<0.05 vs MR by two-tailed Student’s t-test. C, Synergic effect of MR and 5-FU were evaluated by cell counting over 3 days. 3×104 cells were plated in 24-well plate. Relative cell numbers were evaluated by cell counting every 24h. Data are mean ± SEM. * p<0.05 by two-tailed Student’s t-test. D, Rescue effect of Hcy+B12, nucleosides, and NAC alone or in combination on MR and 5-FU-mediated inhibition of cell proliferation by MTT assay. Data are mean ± SEM. * p<0.05 vs control; ^ p<0.05 vs MR; # p<0.05 vs 5-FU; $ p<0.05 vs MR+5-FU by two-tailed Student’s t-test. E, Schematic of reactions from U-^13^C serine tracing study. F, CRC119 and HCT116 cell lines were cultured in the presence (control) or the absence (MR) of methionine with or without addition of 5-FU (3.4 or 10 μM) for 24 h, and then traced with U-^13^C-L-Serine for 6 h. Mass intensity for M+1 dTTP and M+1 methionine. Data are mean ± SD. *p<0.05 compared to the control group and #p<0.05 compared to the MR group by two-tailed Student’s t-test.

Furthermore, we evaluated the combination of MR and 5-FU on cell proliferation in cell culture. We saw a synergistic effect with MR and 5-FU treatments in both primary CRC119 cells and HCT116 cells (Figure 5C). We then conducted similar rescue experiments as described to see which metabolite(s) or metabolic pathways is/are essential for such synergetic effects. Addition of (homocysteine and B12), NAC and formate alone showed no effect on the inhibition of cell proliferation in both lines, whereas nucleosides alone or in combination with (homocysteine+B12) and NAC rescued the MR and 5-FU mediated inhibition of cell proliferation (Figure 5D), highlighting the impairment to nucleotide metabolism. Moreover, we conducted isotope tracing experiments with [U-^13^C]-L-Serine to look further into the MR-induced interruption of nucleotide metabolism (Figure 5E). As shown *in vivo*, MR led to reduction of M+1 dTTP in both CRC119 and HCT116 cell lines. Further combination with 5FU resulted in an additive reduction in M+1 dTTP in both cell lines (Figure 5F). This reduction corresponded with an increase of M+1 methionine (Figure 5F). Thus, the synergy effect between MR and 5FU is at least partially due to an increase in the synthesis of methionine, competing with dTMP synthesis for the availability of serine-derived one carbon unit, 5,10-methylene-THF.

### Dietary MR sensitizes autochthonous Ras-driven sarcoma mouse models to radiation

To explore the therapeutic potential of dietary MR and related mechanisms, we also used a genetically engineered mouse model (GEMM) of soft tissue sarcoma ^40^. GEMMs enable the study of autochthonous cancer development and response to treatment in an anatomically restricted manner ^41^. The two systems together span the spectrum of currently considered pre-clinical tumor models for treatment (i.e. chemotherapy in a PDX for CRC and radiation in an autochthonous model for the sarcoma). Using a site-specific recombinase system (Flp-FRT) ^42^, high-grade sarcomas can be initiated through intramuscular delivery of an adenovirus expressing FlpO recombinase in mice harboring conditional mutations in Kras and Trp53. In brief, extremity sarcomas were induced in *FSF*-*Kras*^*G12D*/+^; *p53*^*FRT*/*FRT*^*(KP*^*FRT*^) mice within 2-3 months after intramuscular delivery of adenovirus expressing FlpO recombinase (Figure 6A). Due to the long duration of tumor onset, we considered the “treatment” study where mice were subjected to dietary treatment when the tumors became palpable (~150 mm^3^). Tumor growth was not altered by dietary treatment alone in this aggressive model (Figure 6B), which is accompanied with minimal effects of MR on cysteine and methionine metabolism in tumors (Figure S5A-B). For soft tissue sarcomas, radiation therapy is often utilized in addition to surgery to achieve local control and can provide a relatively effective treatment option in clinical settings when patients are medically inoperable or refuse surgery ^43^. Thus, we setup to determine the therapeutic possibility of dietary MR in combination with radiation therapy, a dose of 20 Gy focal radiation that is moderately effective in this model ^44^. Strikingly, MR in combination with radiation substantially reduced tumor growth and extended the tumor tripling time by 52%, from an average of 17.48 days for mice on control diet plus radiation to 26.57 days (Figure 6C). Such synergy is comparable to effects seen with known radiosensitizing agents such as pharmacological inhibition of ATM and PI3K/mTOR ^45^. Together, MR sensitizes autochthonous sarcomas with Kras^*G12D*^ mutation and p53 deletion to radiation.

**Figure 6.**
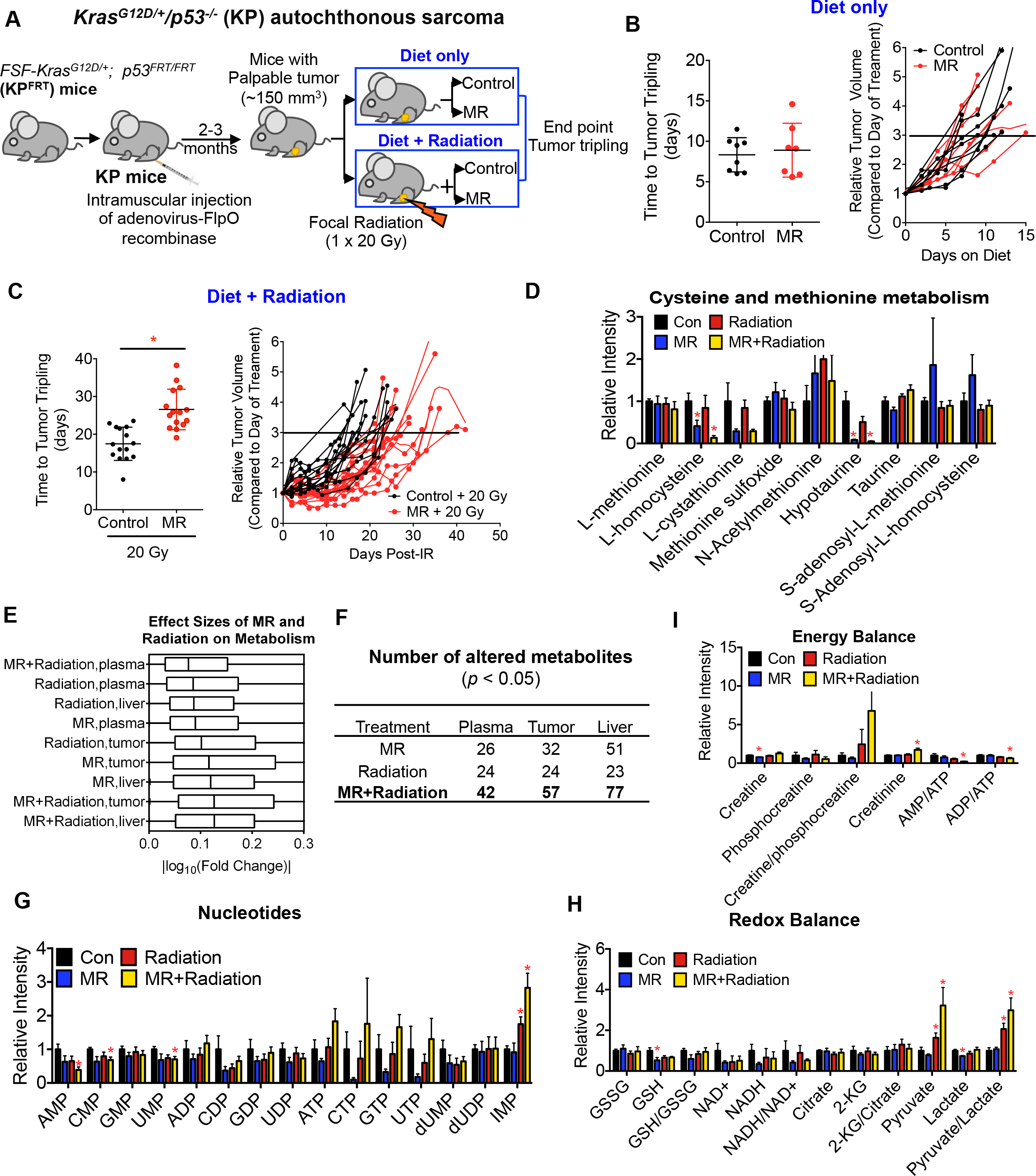
Dietary MR sensitizes RAS-driven autochthonous sarcoma mouse models to radiation. A, Schematic of experimental design. Autochthonous sarcoma model *Kras*^*G12D*/+^/*p53*^−/−^ (KP) mice were used. Soft tissue sarcomas were induced by intramuscular injection of Ad-FlpO into *FSF*-*Kras*^G12D/+^; p53^*FRT*/*FRT*^(KP^FRT^) mice (mixed gender). The onset of tumor in this model took ~2-3 months. When a primary tumor was palpable, mice were randomized to two sets of experiments: in one experiment, mice were subjected to the control or the MR diet when tumor was palpable (Control: n=8, MR: n=7); and in the other experiment, mice were subjected to one focal dose of radiation (20 Gy, Precision X-Ray), followed by the control or the MR diet on the same day until the end point (Control: n=15, MR: n=15). B, Time to tumor tripling and tumor growth curve in mice on dietary treatment only. Data are represented as mean ± SD. C, Time to tumor tripling and tumor growth curve in mice on the combination of dietary treatment and radiation. Data are represented as mean ± SD.*p<0.05. D, Relative intensity of metabolites related to cysteine and methionine metabolism in tumors. Data are mean ± SD.*p<0.05 compared to the control group by two-tailed Student’s t-test. E, Effect of MR, RC (radiation alone), and RM (MR+radiation) on metabolites in tumor, plasma and liver from sarcoma models evaluated by the log_10_ (Fold change). F, Numbers of metabolites significantly changed (*p<0.05, two-tailed Student’s t-test) by MR, RC or RM in plasma, tumor and liver in sarcoma models. G, Relative intensity of nucleotides in tumors. Data are mean ± SD. *p<0.05 compared to the control group by two-tailed Student’s t-test. H, Relative intensity of metabolites related to redox balance and the ratio of GSH/GSSG, NADH/NAD+ and α-KG/Citrate in tumors. GSH, glutathione; GSSG, the oxidized form of glutathione; α-KG, α-ketoglutarate. Data are mean ± SD. *p<0.05 compared to the control group by two-tailed Student’s t-test. I, Relative intensity of metabolites related to energy balance and the ratio of AMP/ATP, ADP/ATP and creatine/phosphocreatine in tumors. Data are mean ± SD.*p<0.05 compared to the control group by two-tailed Student’s t-test.

Metabolomics analysis showed cysteine and methionine metabolism the most altered metabolic pathway by the combination of MR and radiation treatment in liver and plasma, and among the top impacted pathway in tumor as well (Figure S5C-D), although neither methionine or SAM was reduced (Figure 6D), suggesting the synergistic effect might not be related to a reduction in protein synthesis or altered methylation reactions. A similar integrated analysis of the overall changes across the metabolic network in this separate therapeutic context showed the highest correlation (Spearman’s rho = 0.37, p = 2.86e-10) between plasma and liver with alterations in tumor metabolism exhibiting distinct changes (Figure S5C). The largest effects on metabolism occurred in the combination of diet and radiation (Figure 6E-F). In particular, MR in combination with radiation led to a reduction of AMP, CMP, UMP but an increase of IMP, indicating alterations in nucleotide balance (Figure 6G). MR together with radiation also resulted in alterations in cellular redox balance, reflected by a trend towards lower levels of GSH and NADH, and an over 2-fold increase in pyruvate (Figure 6H). Interestingly, MR in combination with radiation also largely decreased the ratios of AMP/ATP and ADP/ATP (Figure 6I), indicating an altered energy metabolism. Meanwhile, the ratio of creatine/phosphocreatine was increased, whereas the creatinine, the waste product of phosphocreatine catabolism, was reduced (Figure 6I). Thus in the sarcoma model, at least three metabolic mechanisms, nucleotide metabolism, energy balance and redox balance, appear to be relevant to the observed synergy of MR and radiation.

### Dietary MR can be achieved in humans

Thus far, we have shown that dietary MR altered systemic cysteine and methionine metabolism in both normal and tumor-bearing laboratory mice. We then questioned whether a similar dietary intervention in healthy humans could achieve a comparable effect on metabolism. In a proof-of-principle clinical study, 6 healthy middle-aged individuals were recruited and subjected to a low methionine (methionine-restricted, MR) diet (methods) for three weeks. The customized MR diet was equivalent to ~2.92 mg/kg daily methionine intake, which resulted in an 83% of reduction in dietary methionine intake (~17.2 mg/kg/d in the baseline level) (Figures 6A-B). As in C57BL/6J mice, dietary MR reproducibly suppressed methionine levels in human subjects (Figures 6C and S6). MR altered circulating metabolism in humans, and cysteine and methionine metabolism remained among the top altered metabolic pathways in plasma (Figure 7D-E). MR reduced N-acetyl-L-cysteine and GSH in all subjects, and altered other metabolites in cysteine and methionine metabolism (Figure 7E). It also affected numerous other metabolites related to methylation, nucleotide metabolism, tricarboxylic acid cycle, and amino acid metabolism (Figures 6F and S6B). Thus, dietary MR affects circulating methionine and cysteine metabolism in healthy humans.

**Figure 7.**
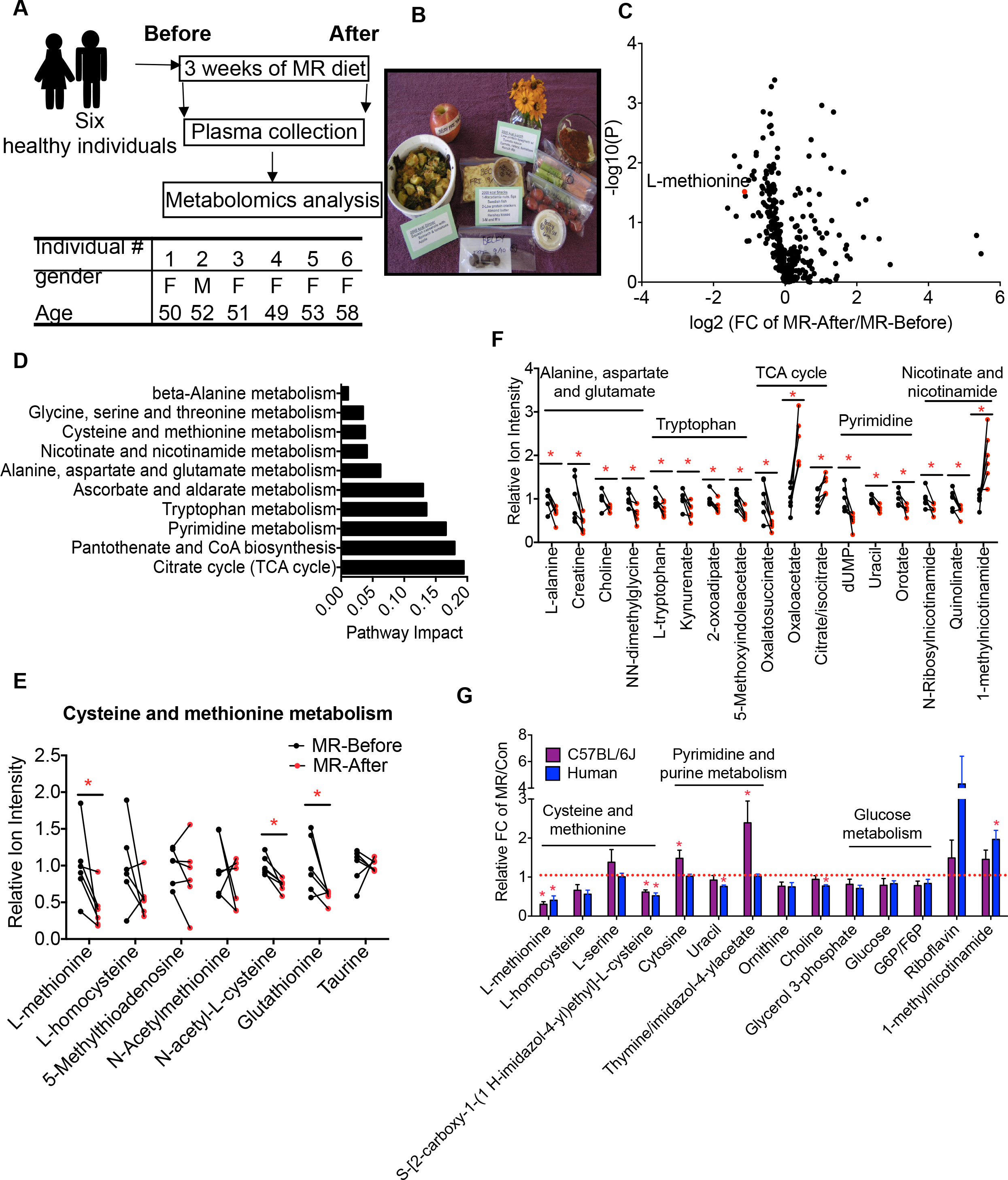
Dietary MR can be achieved in humans. A, Schematic of experimental design, and background information on individuals participated in the dietary study. B, Representative daily MR diet for one individual. C, Volcano plot of metabolites in plasma. D, Pathway analysis of significantly changed (*p<0.05, two-tailed Student’s t-test) plasma metabolites by dietary intervention. E, Relative intensity of plasma metabolites related to cysteine and methionine metabolism. F, Relative intensity of plasma metabolites in the top impacted pathways in D. *p<0.05. G, MR-induced fold changes (FC) of plasma metabolites in cysteine and methionine metabolism, pyrimidine and purine metabolism, and glucose metabolism in non-tumor bearing C57BL/6J mice and healthy human subjects.

We have shown that dietary MR altered circulating metabolites, in particular cysteine and methionine metabolism, in non-tumor bearing C57BL/6J mice (Figure 1), in CRC PDX models (Figures 2–4), in autochthonous sarcomas (Figure 6), and also in healthy human subjects (Figure 7). To integrate the metabolic responses and relate them to tumor physiology, we evaluated the MR-induced changes in metabolites in plasma across the above mouse models and healthy humans. We considered again metabolites related to methionine metabolism as defined earlier in Figure 3F. The Spearman correlation coefficients of the MR response across the different models were highly correlated with the values between 0.53 and 0.73 (Figure S7A-B). Strikingly, the FC of plasma methionine-related metabolites in healthy humans were highly correlated with that in C57BL/6J mice (Spearman’s rho = 0.53, p = 0.1), in the CRC119 mouse model (Spearman’s rho = 0.73, p = 0.02), in the CRC240 mouse model (Spearman’s rho = 0.70, p = 0.02), and in the autochthonous sarcoma model (Spearman’s rho = 0.65, p = 0.04) (Figure S7A-B). Taken together, these results indicate that a conserved response to MR is obtained in several mouse models and in humans. The highest correlations were observed between healthy humans and CRC PDX models. Notably, the metabolite profile in cysteine and methionine metabolism was similar in C57BL/6J mice and in humans (Figure 7G), even though the treatment duration was not the same in the two species. Although dietary MR globally affects metabolism in a specie- and tumor-type- dependent manner, these data show that MR exerts a common effect on methionine-related metabolism that is conserved across mice and humans.

We provide evidence that dietary MR induces a specific metabolic profile involving metabolites related to methionine metabolism in healthy mice and humans (Figure S7D). This controlled clinical study extends observations in methionine free diets that are toxic ^46,47^ to MR at levels that are well tolerated in humans and provide reasonable dietary possibilities including what is possibly obtained in vegan or possibly some pescatarian diets. Future studies assessing how short MR could result in such specific metabolic changes in humans are warranted. These systemic metabolic alterations propagate to changes in cell autonomous one carbon metabolism and lead to responses to MR in a diverse set of pre-clinical models including RAS-driven cancers, including CRC PDXs and autochthonous RAS-driven soft tissue sarcomas. The combined dietary and conventional treatment modalities that target one carbon metabolism result in a cooperative disruption to the flux backbone in one carbon metabolism that creates toxicities involving redox and nucleotide metabolism (Figure S7C). Together this study provides further definitive evidence that dietary alterations that extend beyond the shifting of overall macronutrient balance can have specific effects on the cellular metabolism in distant tissues that is also a component of disease etiology. This work thus supports the intriguing concept of using specific dietary manipulations to selectively target cancer cell metabolism in manners that complement what can be obtained by using radiation or pharmacology, especially in combination settings where the diet and the conventional therapy both target the same pathway such as what we found in one carbon metabolism.

## METHODS

### Animals, diets, and tissue harvesting

All animal procedures and studies were approved by the Institutional Animal Care and Use Committee (IACUC) at Duke University. All mice were housed at 20 ± 2°C with 50 ± 10% relative humidity and a standard12 h dark-12 h light cycle. The special diets with defined methionine levels used previously ^9,17^ were purchased from Research Diets (New Brunswick, NJ, USA) with the control diet contained 0.86% methionine (w/w, catalog #: A11051302) and MR diet contained 0.12% methionine (w/w, catalog #: A11051301). Three mouse models were employed and described below.

### Methionine restriction time course in healthy mice

12-week-old male C57BL/6J mice (Jackson Laboratories, Bar Harbor, ME, USA) were subjected to either the control or the MR diet *ad libitum* for 3 weeks. Mouse blood was sampled through tail bleeding in the morning (10:00 am-12:00 am) at days 1, 2, 4, 7, 10, 14, 17, 21 post-dietary treatments. By day 21, all mice were sacrificed for tissue collection.

### Colorectal cancer PDX models

PDXs of colorectal cancer with liver metastasis were developed as described previously ^34,35^ under an IRB approved protocol (Pro00002435). Briefly, CRC119 and CRC240 tumors were resected, washed and minced as before, and then passaged through JAX NOD.CB17-PrkdcSCID-J mice for 2-5 times. For all the dietary studies, CRC119 and CRC240 PDX tumors were minced in PBS at 150 mg/ml and 200 μl of tumor suspension were subcutaneously injected into the flanks of NOD.Cg-Prkdc^scid^ Il2rg^tm1Wjl^/SzJ mice from the Jackson Laboratory. Mice (3-4 female + 3-4 male) were subjected to the control or the MR diet either two weeks prior to the tumor injection or when tumor was palpable until the end point. Tumor size was monitored two to three times per week until the end point when tumor reached ~1,000 mm^3^. For the combination therapy with a standard chemo drug 5-FU, mice were subjected to the control or the MR diet two weeks prior to the tumor injection until the end of the study when tumor size reached ~1,000 mm^3^. When tumor was palpable, mice (4 female + 4 male) were randomized to treatment of 5-FU (NDC 63323-117-10, 12.5 mg/kg three times per week) or vehicle (saline) through intraperitoneal injection. Tumor size was monitored two to three times per week until the end point when tumor reached ~1,000 mm^3^.

### Autochthonous soft tissue sarcomas

Primary soft-tissue sarcomas were generated as described previously ^40,42^. Briefly, *p53*^*FRT*^ mice were crossed with mice carrying an Flp-activated allele of oncogenic Kras (*FSF*-*Kras*^*G12D*^) to generate FSF-*Kras*^*G12D*/+;^*p53*^*FRT*/*FRT*^(*KP*^*FRT*^) compound conditional mutant mice. *p53*^*FRT*^ mice and *FSF*-*Kras*^*G12D*^ mice were maintained on mixed C57BL/6J × 129SvJ backgrounds. Soft tissue sarcomas were induced by intramuscular injection of Ad-FlpO into *KP*^*FRT*^ mice. 25 μl Ad5CMVFlpO (6 × 10^10^ PFU/ml) was incubated with 600 μl minimum essential media (Sigma-Aldrich, St Louis, MO) and 3 μl 2 M CaCl_2_ (Sigma-Aldrich, St Louis, MO) for 15 minutes to form calcium phosphate precipitates. 50 μl precipitated virus was injected intramuscularly per mouse to generate sarcomas. Soft tissue sarcomas developed at the site of injection in the lower extremity as early as 2 months post injection. *FSF*-*KrasG12D* mice were kindly provided by Tyler Jacks (MIT) and we previously generated p53^FRT^mice at Duke University ^40,42^.

KP mice (mixed gender) were subjected to control or MR diet when tumor was palpable (~150 mm^3^) until the end point when tumor tripled. Tumor size was monitored 2-3 times per week. For combination therapy with radiotherapy, KP mice with palpable tumor were subjected to a single dose of 20 Gy focal irradiation using the X-RAD 225Cx small animal image-guided irradiator (Precision X-Ray). The irradiation field was centered on the target via fluoroscopy with 40 kilovolt peak (kVp), 2.5 mA x-rays using a 2-mm aluminum filter. Sarcomas were irradiated with parallel-opposed anterior and posterior fields with an average dose rate of 300 cGy/min prescribed to midplane with 225 kVp, 13 mA x-rays using a 0.3-mm copper filter and a collimator with a 40 × 40 mm^2^ radiation field at treatment isocenter. The dose rate was monitored in an ion chamber by members of the Radiation Safety Division at Duke University. After radiation, mice were subjected to the control or the MR diet immediately until the end point when tumor tripled. Tissues (tumor, liver and plasma) were collected at the time of tumor tripling. For metabolomics analysis, another cohort of mice on the combination therapy with radiation were sacrificed at 10 days post radiation and dietary treatment, the average time point when the KP mice on the control or the MR diet alone had their tumor size tripled.

### Tissue collection

For tissue collection from all the above animal studies, mice were fasted in the morning for 4 h (9:00-13:00). Tumor, plasma and liver were collected followed by immediate snap frozen, and stored at − 80 °C until processed.

### Colorectal cancer cell lines

Early passage colorectal cancer cell lines, CRC119 and CRC240 cell lines, were developed from the PDXs. PDXs were harvested and homogenized, and the homogenates were grown in RPMI 1640 media with addition of 10% fetal calf serum, 100,000 U/L penicillin and 100 mg/L streptomycin at 5% CO_2_. A single cell clone was isolated using an O ring. Cell lines were authenticated at the Duke University DNA Analysis Facility by analyzing DNA samples from each cell lines for polymorphic short tandem repeat (STR) markers using the GenePrint 10 kit from Promega (Madison, WI, USA). HCT116 cell line was a gift from Dr. Lewis Cantley’s laboratory, and was maintained in RPMI 1640 supplemented with 10% fetal bovine serum and 100,000 U/L penicillin and 100 mg/L streptomycin. Cells were grown at 37 °C with 5% CO_2_.

### Cell viability assay

Cell viability was determined by MTT assay. Briefly, cells cultured at 96-well plates were incubated in RPMI medium containing MTT (final concentration 0.5 mg/ml) at cell incubator for 2-4 hours. Media were then removed and replaced with 100 μl of DMSO, followed by additional 10 min of incubation at 37 °C. The absorbance at 540 nm was read using a plate reader. For the metabolite rescue studies, the following final conditions of metabolites were used: homocysteine (400 μM), B12 (20 μM), nucleosides (1x) choline (1 mM), formate (0.5 mM), NAC (1 mM), 5-FU (2.5 μM).

### Human dietary study

The controlled feeding study was conducted at Penn State University Clinical Research Center (CRC) and approved by the Institutional Review Board of the Penn State College of Medicine in accordance with the Helsinki Declaration of 1975 as revised in 1983 (IRB# 32378). Healthy adults of mixed gender were recruited by fliers and word of mouth and, as assessed for initial eligibility by telephone interview, were free of disease and currently not taking certain medications including anti-inflammatory drugs, corticosteroids, statins, thyroid drugs, and oral contraceptives. Final eligibility was assessed by standard clinical chemistry and hematology analyses. Written consent was obtained from eligible subjects and baseline resting metabolic rate was assessed by indirect calorimetry (Parvo Medics), physical activity by questionnaire and dietary intake by 3 unannounced 24 h diet recalls conducted by telephone in the week prior to returning to the clinic. All subjects were placed on an MR diet for the final 3 weeks which provided 50-53% of energy from carbohydrate, 35-38% from fat, and 12-13% from protein and total calories adjusted individually based upon the baselines of resting metabolic rate and physical activity (calculated by the Harris Benedict Equation). Of the total protein, 75% was provided by a methionine-free medicinal beverage (Hominex-2, Abbott Nutrition, Columbus, OH) and the remaining 25% was from low methionine foods such as fruits, vegetables and refined grains. The total methionine intake was ~2.92 mg//kg/day, representing an 83% reduction in methionine intake compared to pre-test values from diet recalls. The five-day cycle menu was created and evaluated for nutrient content using the Nutrition Data System for Research (2006, Minneapolis, MN). Blood was sampled into EDTA tubes in the morning after overnight fasting by a registered nurse at the beginning and end of the diet period. Plasma was obtained after centrifugation at 5,000 rpm for 10 min at 4°C. All 6 subjects agreed to have their samples and data used for future research.

### Metabolite profiling and isotope tracing

PDX primary cells were seeded in 6-well plates at a density of 2.0 × 10^5^ cells per well. For overall polar metabolite profile, after overnight incubation, cells were washed once with PBS and cultured for an additional 24 h with 2 ml of conditional RPMI medium containing 0.1 μM or 0 μM methionine plus 10% FBS. Cellular metabolites were extracted after incubation. For [U-^13^C]-L-Serine isotope tracing, both CRC119 and HCT116 cell lines HCT116 cells were seeded in 6-well plates at a density of 2.0 × 10^5^ cells per well. Cells were washed once with PBS after overnight incubation and cultured for an additional 24 h with 2 ml of conditional RPMI medium containing 0 μM or 100 μM methionine with or without addition of 5-FU (3.4 or 10 μM) plus 10% FBS. Then, medium was replaced with fresh conditional RPMI medium (0 μM or 100 μM methionine) with or without addition of 5-FU (3.4 or 10 μM) containing tracer U-^13^C]-L-Serine plus 10% dialyzed FBS. Cells were traced for 6 h, and followed by cellular metabolite extraction.

### Metabolite extraction

Polar metabolite extraction has been described previously (Liu et al., 2015). Briefly, tissue samples, liver and tumor, were pulverized in liquid nitrogen and then 3-10 mg of was weighed out for metabolite extraction using ice cold extraction solvent (80% methanol/water, 500 μl). Tissue was then homogenized with a homogenizer to an even suspension, and then incubated on ice for an additional 10 min. The extract was centrifuged at 20 000 g × 10 min at 4 °C. The supernatant was transferred to a new Eppendorf tube and dried in vacuum concentrator. For serum or medium, 20 μl of sample was added to 80 μl ice cold water in an Eppendorf tube on ice, followed by the addition of 400 μl ice cold methanol. Samples were vortexed at the highest speed for 1 min before centrifugation at 20,000 g for 10 min at 4 °C. For cells cultured in 6-well plates, cells were placed on top of dry ice right after medium removal. 1 ml ice-cold extraction solvent (80% methanol/water) was added to each well and the extraction plate was quenched at −80 °C for 10 min. Cells were then scraped off the plate into an Eppendorf tube. Samples were vortexed and centrifuged as described early. The supernatant was transferred to a new Eppendorf tube and dried in vacuum concentrator. The dry pellets were stored at −80 °C for LC-HRMS analysis. Samples were reconstituted into 30-60 μl sample solvent (water:methanol:acetonitrile, 2:1:1, v/v/v) and were centrifuged at 20,000 × *g* at 4 °C for 3 min. The supernatant was transferred to LC vials. The injection volume was 3 μl for HILIC chromatography, which is equivalent to a metabolite extract of 160 μg tissue injected on the column.

### HPLC method

Ultimate 3000 UHPLC (Dionex) was coupled to Q Exactive-Mass spectrometer (QE-MS, Thermo Scientific) for metabolite separation and detection. For additional polar metabolite analysis, a hydrophilic interaction chromatography method (HILIC) with an Xbridge amide column (100 × 2.1 mm i.d., 3.5 μm; Waters) was used for compound separation at room temperature. The mobile phase and gradient information was described previously (Liu et al., 2014).

### Mass spectrometry and data analysis

The QE-MS was equipped with a HESI probe, and the relevant parameters were as listed: heater temperature, 120 °C; sheath gas, 30; auxiliary gas, 10; sweep gas, 3; spray voltage, 3.6 kV for positive mode and 2.5 kV for negative mode. Capillary temperature was set at 320 °C, and S-lens was 55. A full scan range was set at 60 to 900 (m/z) when coupled with the HILIC method, or 300 to 1000 (m/z) when low abundance metabolites need to be measured. The resolution was set at 70 000 (at m/z 200). The maximum injection time (max IT) was 200 ms. Automated gain control (AGC) was targeted at 3 Å~ 10^6^ ions. LC-MS peak extraction and integration were analyzed with commercial available software Sieve 2.0 (Thermo Scientific). The integrated peak intensity was used for further data analysis. For tracing studies using U-^13^C]-L-Serine, ^13^C natural abundance was corrected as previously described ^48^.

### Statistical analysis and bioinformatics

Pathway analysis of metabolites was carried out with software Metaboanalyst (http://www.metaboanalyst.ca/MetaboAnalyst/) using the KEGG pathway database (http://www.genome.jp/kegg/). All data are represented as mean ± SD or mean ± SEM as indicated. *P* values were calculated by a two-tailed Student’s *t* test unless otherwise noted.

### Singular value decomposition analysis of time-course metabolomics data

We firstly constructed a combinational matrix containing raw ion intensities of plasma metabolites from C57BL/6J mice, both the control and the MR groups. For each group there were 9 time points and 5 replicates for each time point, resulting in a 311 × 90 matrix. This matrix was then log-transformed and iteratively row-normalized and column-normalized until mean values of all rows and columns converged to zero. Singular value decomposition was applied on the processed matrix to identify dominating dynamic modes:

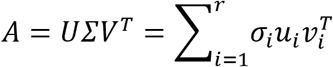

In which A is the processed metabolomics matrix, σ_*i*_ is the ith singular value (ranked from maximal to minimal), 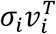 is termed the ith mode. Modes 2 and 3 were defined as responding modes due to significant difference between control and MR values in both modes. For the ith metabolite, total contribution of modes 2 and 3 to its dynamics was evaluated by:

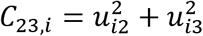

Time-course metabolomics data of 50 metabolites with highest contribution of modes 2 and 3 was then clustered using clustergram() function in MATLAB R2016b. All methods used were implemented in MATLAB codes.

### Cross-tissue comparison of metabolite FC in PDX and sarcoma models

Spearman’s rank correlation coefficients were computed on metabolites measured in the plasma, tumor and liver. Distance between FC in tissues A and B (*e.g.* liver and tumor) was computed by Euclidean distances between the two vectors of FC of all metabolites measured in both A and B. Multidimensional scaling (MDS) was then applied to visualize the tissues in two-dimension, in which a stress function quantifying how accurate the pairwise distances between points are represented in the dimension-reduced coordinates compared to the original dataset was minimized:

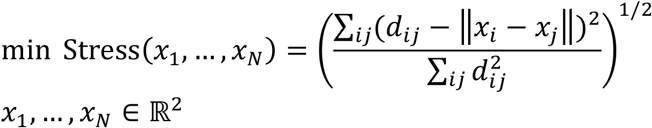

In which *d*_*ij*_ is the distance between the ith and jth data points in the original dataset and *x*_*i*_ means the ith point in the dimension-reduced dataset. All methods used here were implemented in MATLAB codes.

### Methionine-related and -unrelated metabolites

Methionine-related and -unrelated metabolites were defined according to their distance to methionine in the genome-scale metabolic model of human, Recon 2 ^36^. Metabolites were defined as methionine-related if distance to methionine is less than or equal to 4. All metabolites with distance to methionine larger than 4 were defined as methionine-unrelated. Metabolites were mapped by their KEGG IDs between the metabolomics dataset and Recon 2.

### Quantitation of methionine concentrations

To quantify methionine concentrations in the plasma, liver and tumor across the mouse models and in healthy humans, two additional datasets of metabolomics profiles in human plasma with their corresponding absolute methionine concentrations quantified using ^13^C-labeled standards were used. The raw intensities across all samples were log-transformed and normalized. Linear regression was then performed on the normalized datasets to predict absolute methionine concentrations. Four normalization algorithms including cyclic loess, quantile, median and z-score were tested. Among the normalization algorithms, cyclic loess had the highest R-squared statistics in the corresponding linear regression model (R^2^=0.74 for cyclic loess compared to 0.66 for quantile, 0.68 for median and 0.70 for z-score). Thus, the cyclic loess normalized dataset was used for the final model training, which generated the following equation describing the model:

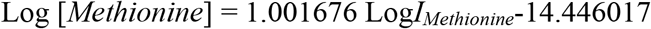

In this equation, [*Methionine*] is the absolute methionine concentration, *I*_*Methionine*_ is the cyclic loess normalized value of methionine intensity.

### Data and Software Availability

The metabolomics data reported in this study were deposited to Mendeley Data (DOI: doi:10.17632/zs269d9fvb.1).

## AUTHOR CONTRIBUTIONS

X.G. and J.W.L. designed the study, X.G., and J.W.L. wrote and edited the paper. D.E.C. and D.G.K designed the sarcoma experiments and edited the paper. M.L. and D.S.H designed and implemented the colorectal PDX models and edited the paper. X.G., M.L, D.E.C., G.A., and M.A.R. performed animal experiments. X.G., S.M.S, and M.A.R performed all cell culture experiments. J.P.R., A.C., A.C., and S.N.N. conducted the human study. Z.D. conducted computational analyses with initial help from P.M. J.L. and M.A.R assisted in mass spectrometry metabolomics experiments.

## ACKNOWLEDGEMENTS

Support from the National Institutes of Health R01CA193256, R21CA201963, P30CA014236 (J.W.L.), R35CA197616 (D.G.K.), T32CA93240 (D.E.C.), and a fellowship from the Canadian Institutes of Health Research (CIHR) (X.G., funding reference number 146818) are gratefully acknowledged. We thank Drs. Mary Lou Kiel and Terryl Hartman for their assistance in designing the diets and Sami Heim for her help with food preparation in the human study. The human study was partially supported by the CRC at Penn State University (NIH M01 RR 10732). We gratefully acknowledge members of the Locasale lab for helpful discussions.

## DISCLOSURES

A provisional patent related to this work has been filed.

---

**Figure S1.**
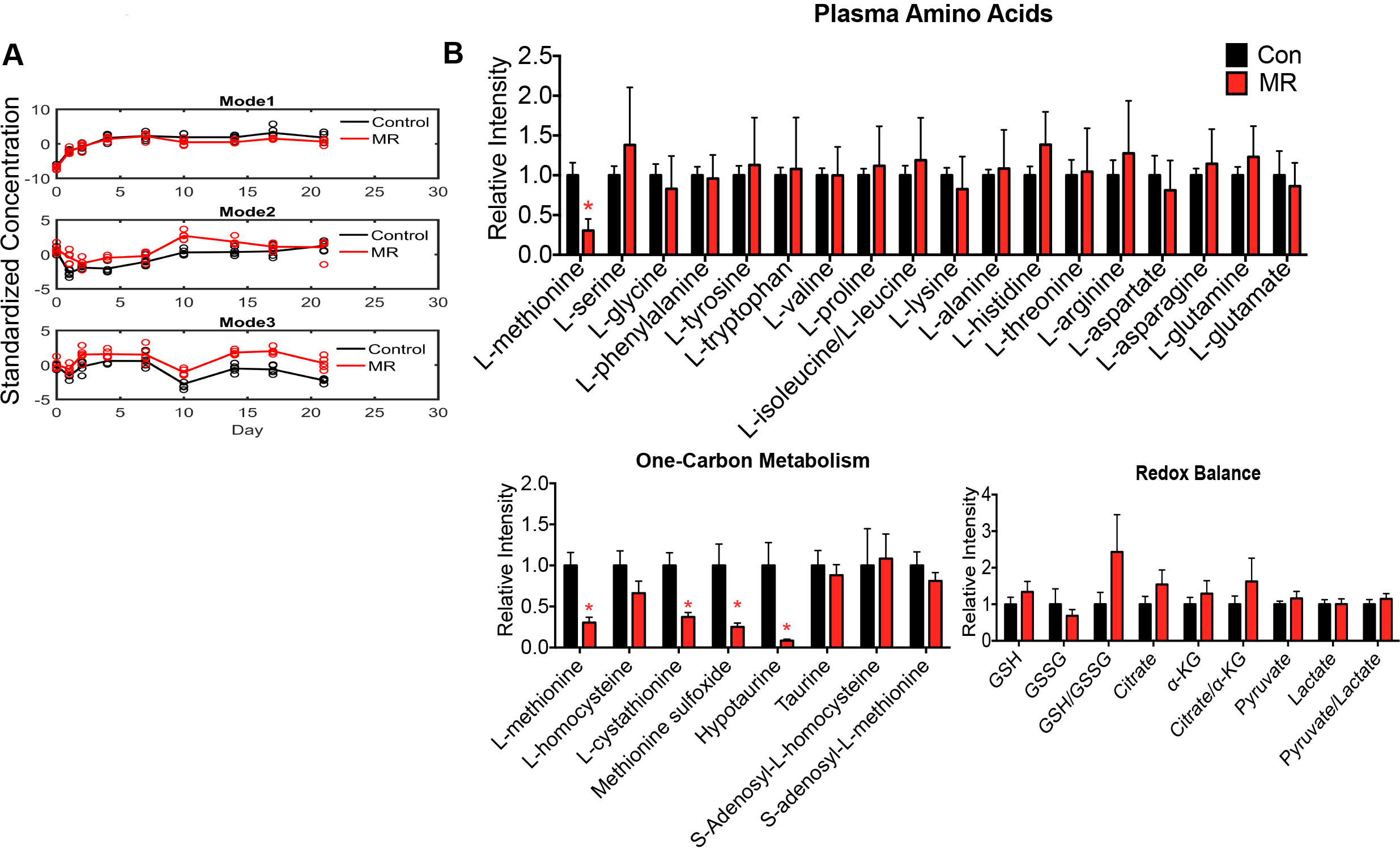
Related to Figure 1. A, Dynamic patterns of top 3 modes. Standardized concentration (the values are normalized to have mean=0, standard deviation=1) in Mode 1, mode 2 and mode 3. B, Relative intensity of amino acids in plasma collected at the end of study. Data are represented as mean ± SD.*p<0.05, two-tailed Student’s t-test. C-D, Metabolomics analysis of plasma metabolites from both female and male C57BL/6J mice subjected to control and MR diet for 12 weeks. C, Volcano plots of metabolites in female and male mice. D, Metabolites related to methionine metabolism such as homocysteine, taurine and S-adenosyl-methionine in both male and female. Data are represented as mean ± SD. *p<0.05, two-tailed Student’s t-test. E, Pathway analysis of significantly changed (p<0.05 by two tailed student t-test) metabolites by MR.

**Figure S2.**
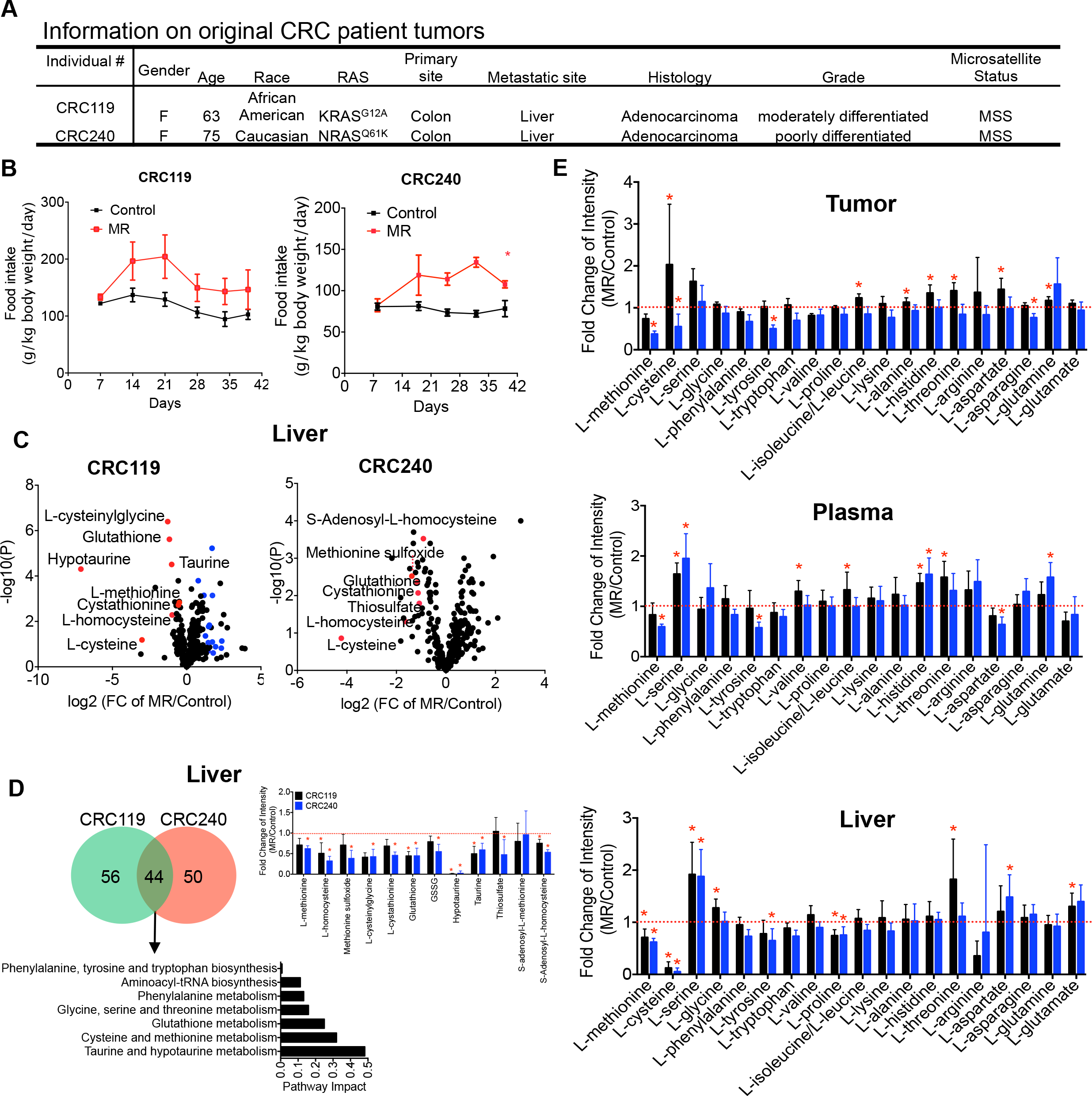
Related to Figure 2. A, Information on original CRC patient tumors. B, Food intake in CRC119 and CRC240 models in Figure 2C.

**Figure S3.**
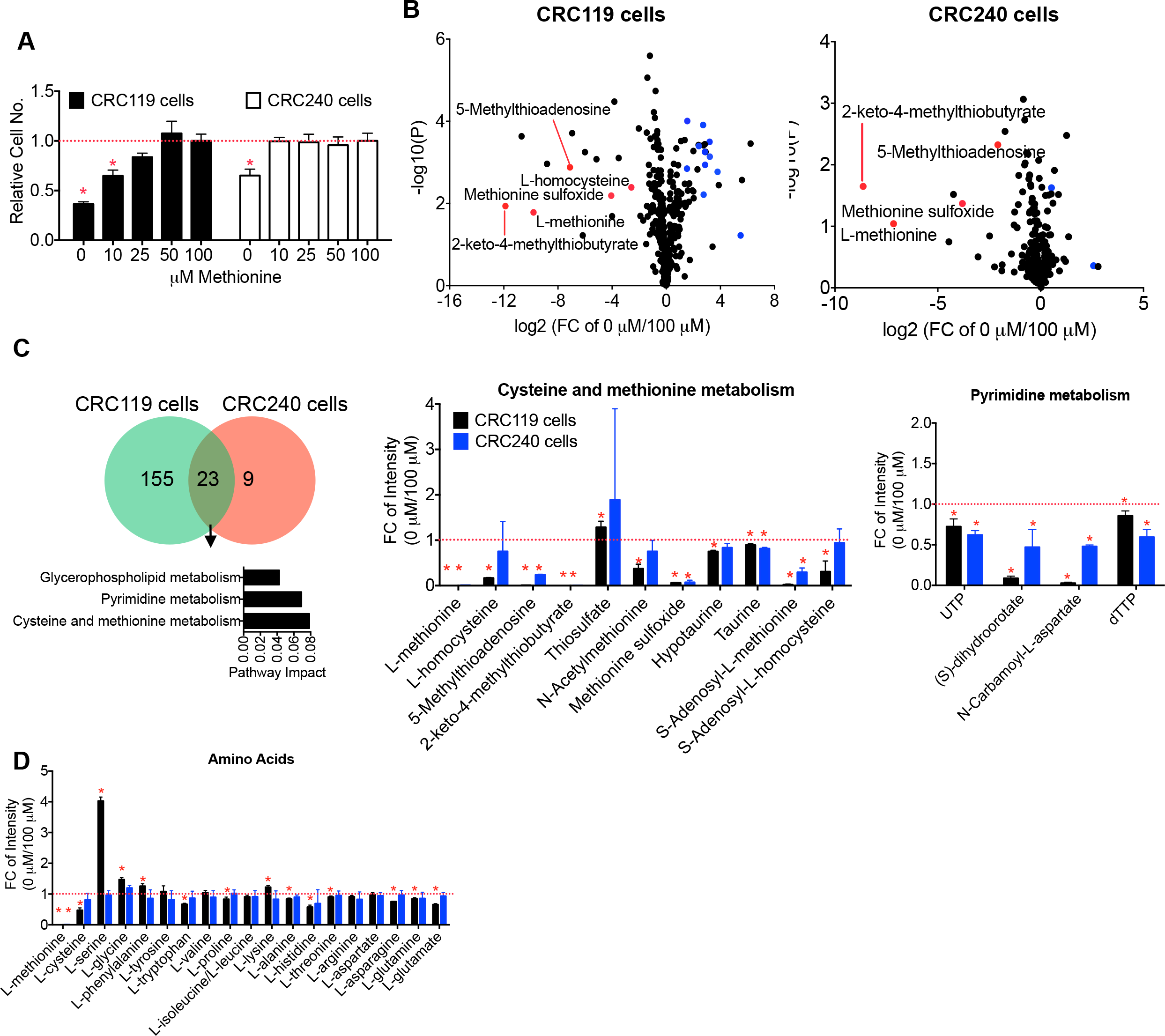
Related to Figure 3. A, Relative cell numbers in CRC119 and CRC240 primary tumor cells treated with different doses of methionine for 72 h. B, Volcano plots of metabolites in cells treated with 0 or 100 μM methionine for 24 h. C, Left: Venn diagram of changed metabolites in CRC119 and CRC240 primary tumor cells treated with 0 or 100 μM methionine, and pathway analysis of commonly changed metabolites. Right: Fold changes (FC) of metabolites in the cysteine and methionine metabolism and pyrimidine metabolism in CRC119 and CRC240 primary tumor cells treated with 0 or 100 μM methionine. D, Relative FC of intensity of amino acids by methionine deprivation in CRC119 and CRC240 primary tumor cells. Data are represented as mean ± SD.*p<0.05, two-tailed Student’s t-test.

**Figure S4.**
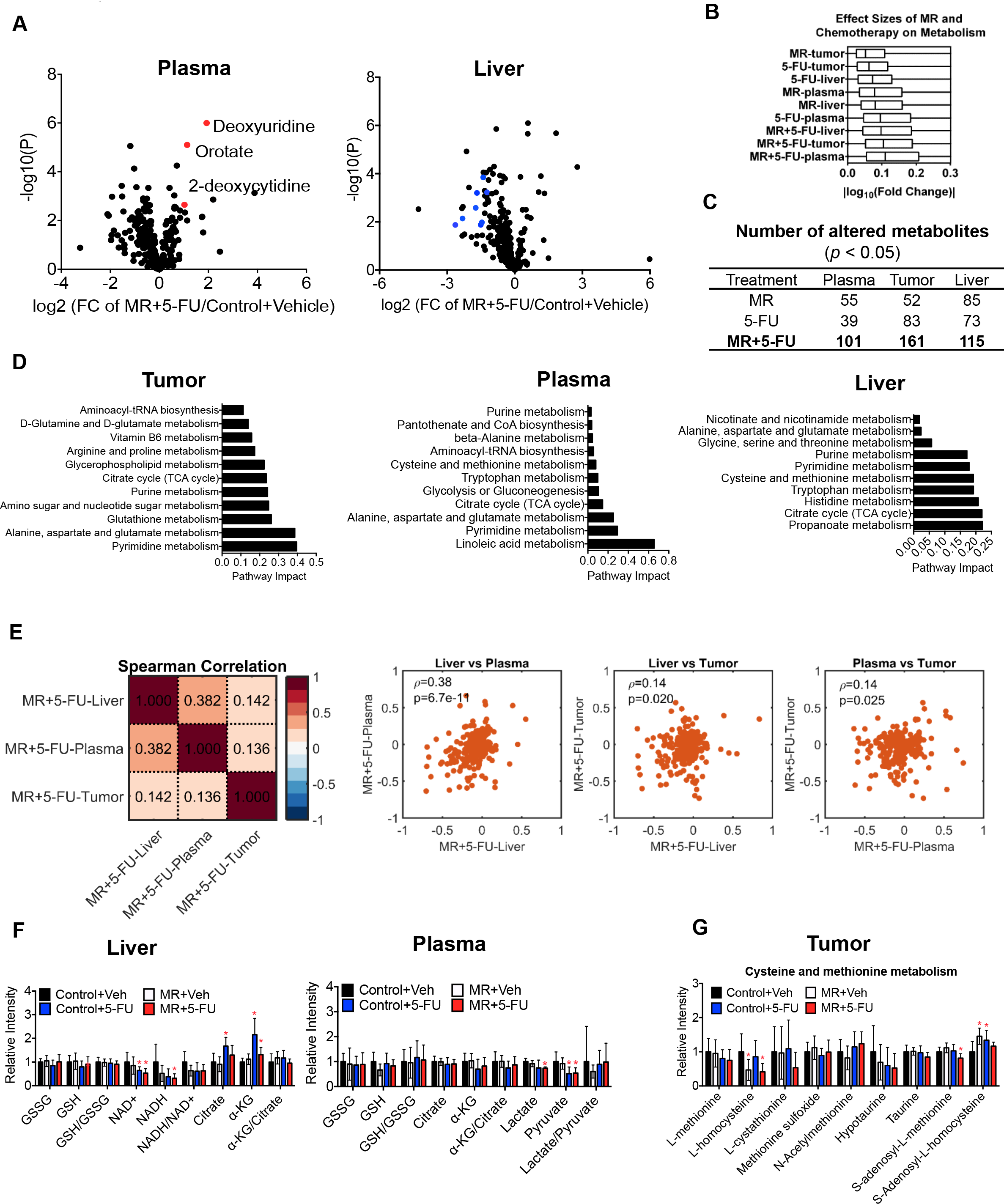
Related to Figure 4. A, Volcano plots of metabolites in plasma and liver altered by the combination of dietary MR and 5-FU. FC, fold change. Blue dots indicate acylcarnitines. B, Effect of 5-FU alone and a combination of MR and 5-FU on metabolites in tumor, plasma and liver from sarcoma models, evaluated by the log_10_ (Fold change). C, Numbers of metabolites significantly changed (*p<0.05, two-tailed Student’s t-test) by MR, 5-FU or a combination of MR and 5-FU in plasma, tumor and liver. D, Pathway analysis (false discovery rate < 0.5) of metabolites significantly changed (*p<0.05, two-tailed Student’s t-test) by MR, 5-FU, or by the combination of dietary MR and 5-FU. E, Spearman’s rank correlation coefficients of MR and 5-FU -induced fold changes of metabolites in tumor, plasma and liver from mice on dietary MR and 5-FU. F, Relative intensity of metabolites related to redox balance and the ratio of GSH/GSSG, NADH/NAD+ and α–KG/Citrate or lactate/pyruvate in the liver and plasma. GSH, glutathione; GSSG, the oxidized form of glutathione; α–KG, α–ketoglutarate. G, Relative intensity of metabolites related to cysteine and methionine metabolism.

**Figure S5.**
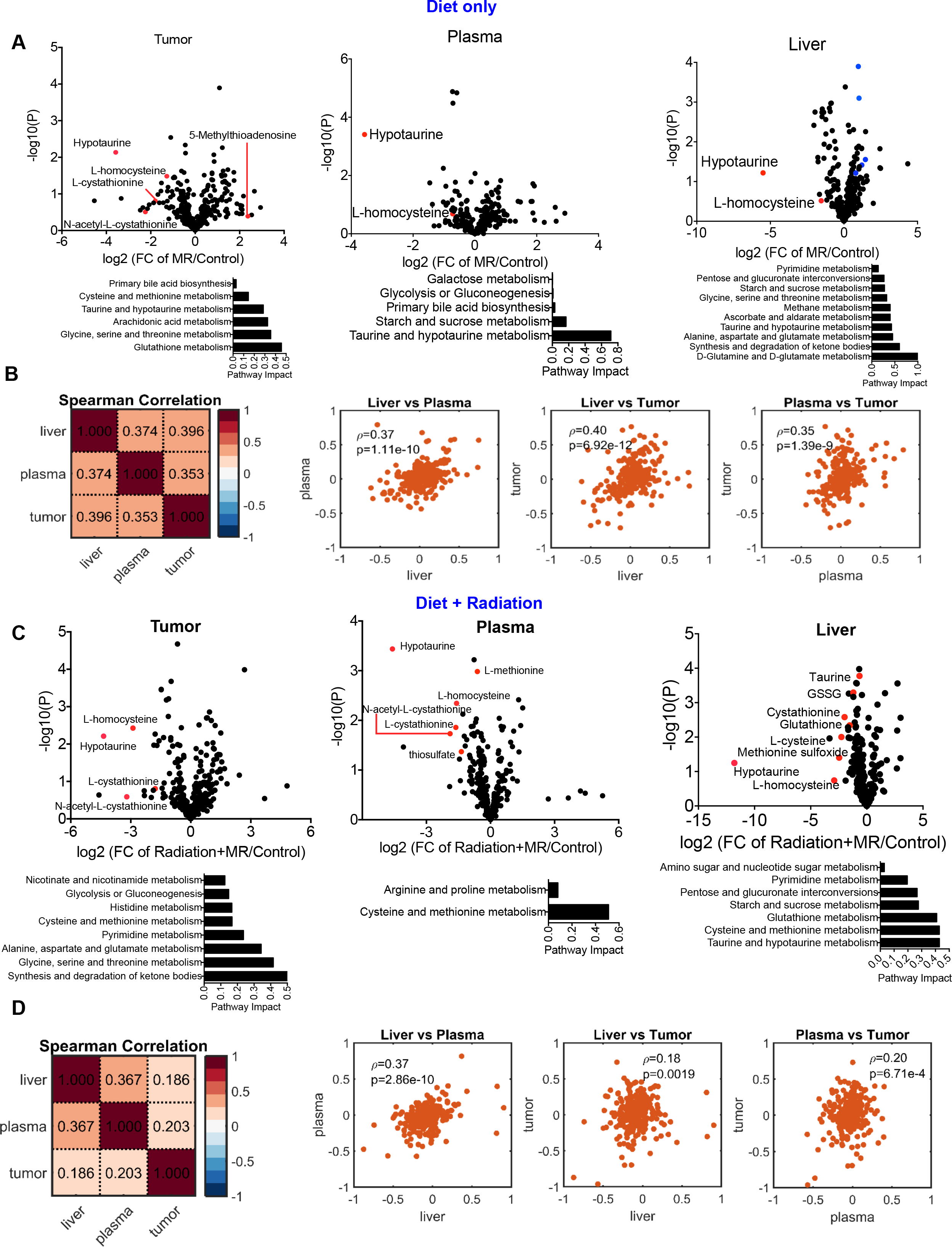
Related to Figure 6. A, Volcano plots of metabolites in tumor, plasma and liver, and pathway analysis of metabolites significantly changed (*p<0.05, two-tailed Student’s t-test) by dietary MR alone. Blue dots indicate acylcarnitines. B, Spearman’s rank correlation coefficients of MR-induced fold changes of metabolites in tumor, plasma and liver from mice on dietary treatment alone. C, Volcano plots of metabolites in tumor, plasma and liver, and pathway analysis (false discovery rate < 0.5) of metabolites significantly changed (*p<0.05, two-tailed Student’s t-test) by dietary MR and radiation. Blue dots indicate acylcarnitines. D, Spearman’s rank correlation coefficients of MR and radiation-induced fold changes of metabolites in tumor, plasma and liver from mice on dietary MR and radiation.

**Figure S6.**
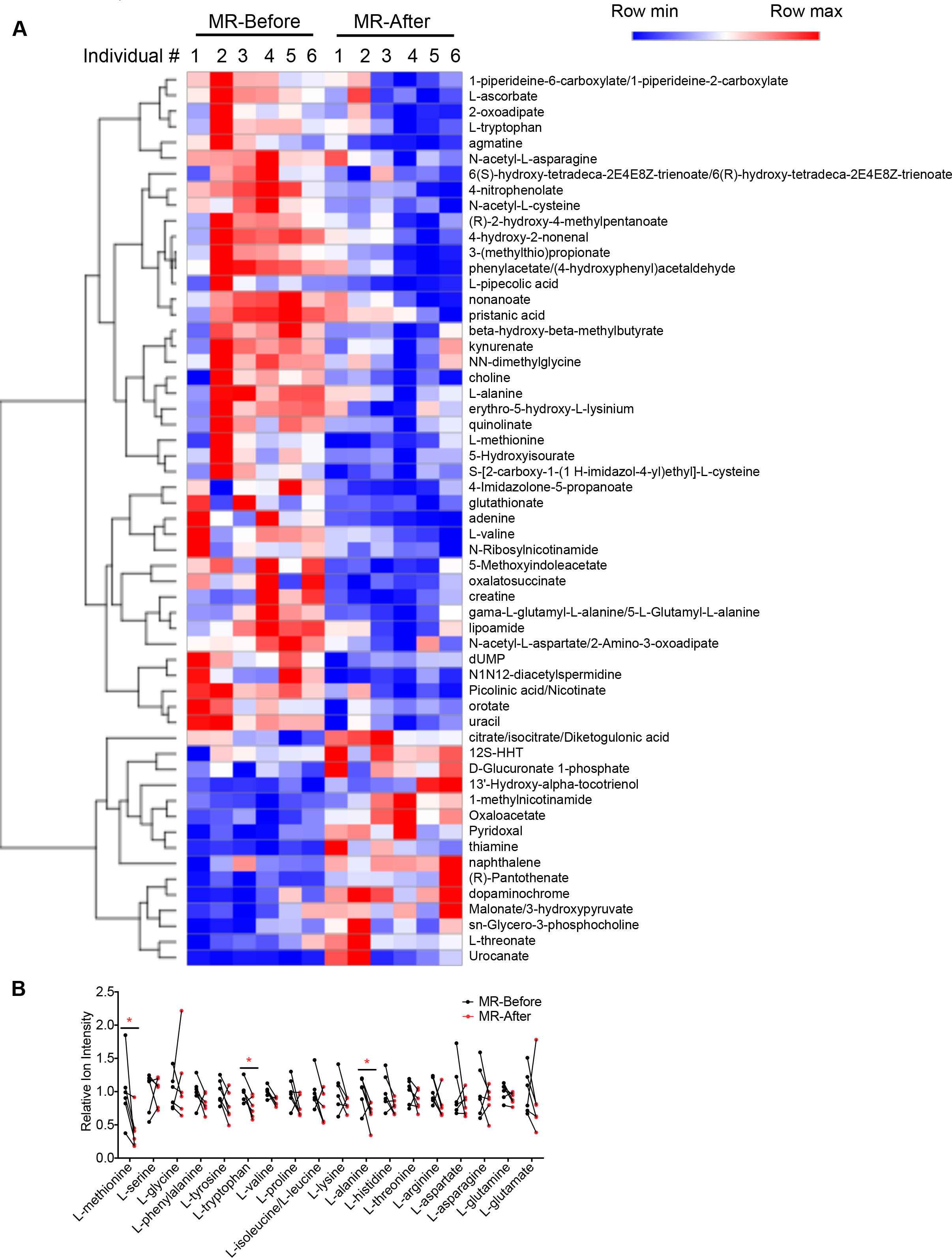
Related to Figure 7. A, Heatmap of significantly changed (*p<0.05, two-tailed Student’s t-test) plasma metabolites by dietary intervention in human subjects. B, Relative intensity of amino acids in plasma. *p<0.05.

**Figure S7.**
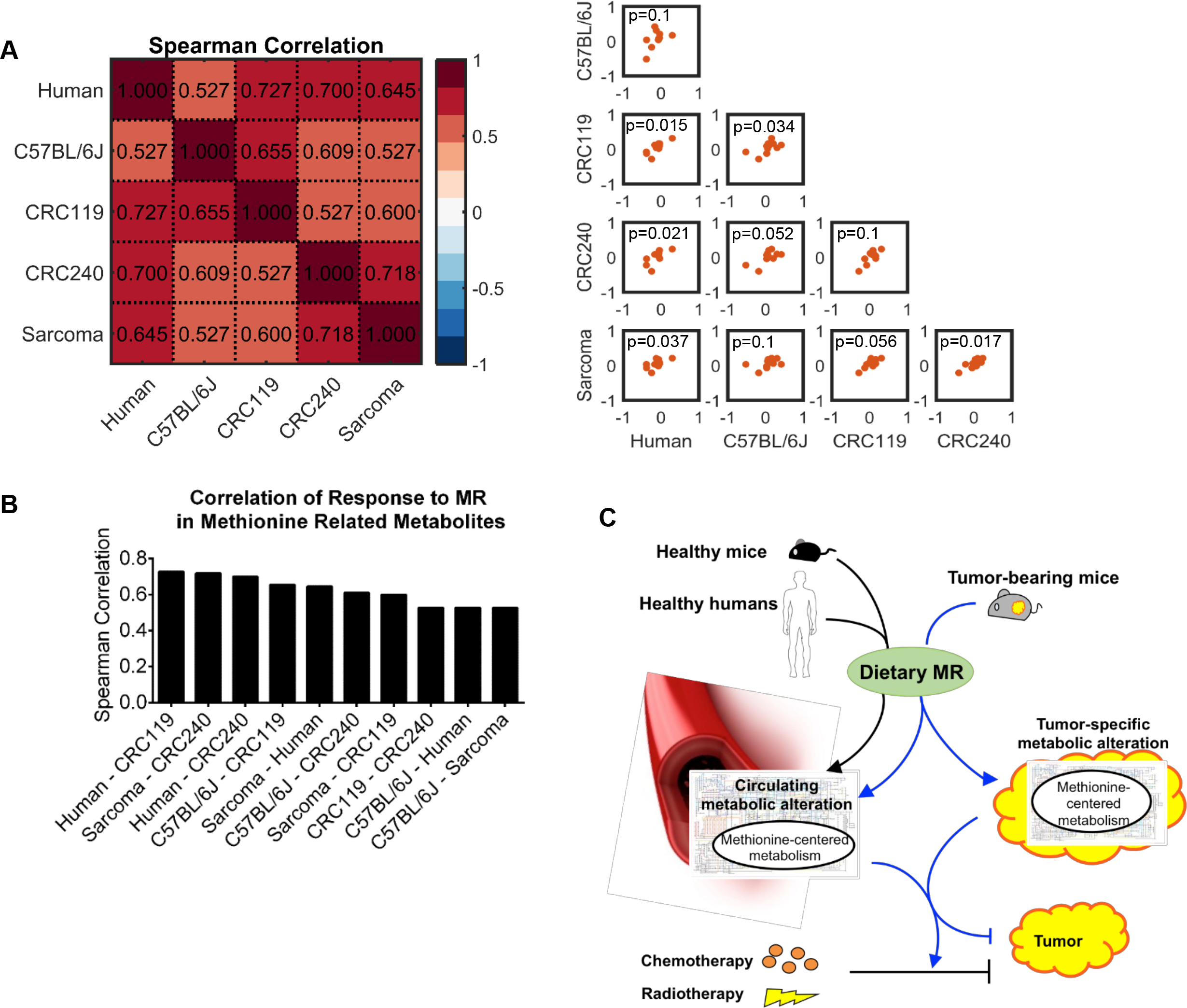
Comparative metabolic effects of MR across mouse models and humans. A, Spearman’s rank correlation coefficients of MR-induced fold changes of methionine-related metabolites (defined in Figure 3F) in plasma samples from non-tumor bearing C57BL/6J mice, PDX CRC119 and CRC240 mouse models, sarcoma mouse model, and healthy human subjects. B, Spearman’s rank correlation coefficients among different models in A ranked from the highest to the lowest. C, Model: MR alters circulating metabolites centered to methionine metabolism in health mice and human. Such alteration correlates with the changes in tumor-bearing mouse models, and synergizes with chemo- and radio-therapy. Data are represented as mean ± SD. *p<0.05.

**Table S1.** Methionine concentrations in the plasma, tumor and liver across mouse models and humans. Tissues were collected at the end of each experiment. Concentrations in tissues were estimated in μM, assuming 1 g wet tissue weight = 1 ml. Quantitation was performed by using ^13^C-labeled standards for each amino acid, which were added before extraction. Cyclic Loess normalization and linear regression was applied in quantitation of methionine in samples without ^13^C-labeled standards. Values are mean ± SD. *p < 0.05 by Student t-test between the control diet and the MR diet, #p < 0.05 by Student t-test between the control diet vs the MR diet + radiation.

| Model | Tissue | Control diet | MR diet |
| --- | --- | --- | --- |
| CRC119 | Plasma | 37.58 ± 6.08 | 27.44 ± 4.28* |
| | Tumor | 31.23 ± 9.10 | 20.08 ± 2.36* |
| | Liver | 29.99 ± 2.38 | 18.68 ± 4.12* |
| CRC240 | Plasma | 52.35 ± 12.19 | 32.98 ± 3.60* |
| | Tumor | 70.77 ± 25.73 | 31.82 ± 3.05* |
| | Liver | 18.97 ± 3.10 | 12.393 ± 1.86* |
| CRC119 (Vehicle) | Plasma | 35.93 ± 6.02 | 33.88 ± 7.97 |
| | Tumor | 33.52 ± 10.31 | 31.50 ± 12.12 |
| | Liver | 17.37 ± 4.18 | 16.68 ± 3.88 |
| CRC119 (5-FU) | Plasma | 32.13 ± 5.29 | 31.31 ± 11.55 |
| | Tumor | 26.70 ± 7.15 | 29.45 ± 11.04 |
| | Liver | 16.50 ± 5.14 | 16.61 ± 2.39 |
| Sarcoma | Plasma | 37.75 ± 5.27 | 35.86 ± 14.57 |
| | Tumor | 46.84 ± 10.01 | 38.39 ± 10.32 |
| | Liver | 10.44 ± 4.74 | 10.45 ± 3.44 |
| Sarcoma (Radiation) | Plasma | 24.44 ± 7.56 | 25.87 ± 4.11# |
| | Tumor | 43.10 ± 11.86 | 37.22 ± 4.97# |
| | Liver | 10.43 ± 2.98 | 6.55 ± 1.9* |
| C57BL/6J | Plasma | 99.89 ± 16.90 | 28.79 ± 4.59* |
| Human | Plasma | 13.74 ± 5.19 | 6.55 ± 4.02* |

## REFERENCES

1 Cantor, J. R. et al. Physiologic Medium Rewires Cellular Metabolism and Reveals Uric Acid as an Endogenous Inhibitor of UMP Synthase. Cell 169, 258–272 e217, doi:10.1016/j.cell.2017.03.023 (2017).

2 Tardito, S. et al. Glutamine synthetase activity fuels nucleotide biosynthesis and supports growth of glutamine-restricted glioblastoma. Nat Cell Biol 17, 1556–1568, doi:10.1038/ncb3272 (2015).

3 Liu, X., Romero, I. L., Litchfield, L. M., Lengyel, E. & Locasale, J. W. Metformin Targets Central Carbon Metabolism and Reveals Mitochondrial Requirements in Human Cancers. Cell Metab 24, 728–739, doi:10.1016/j.cmet.2016.09.005 (2016).

4 Maddocks, O. D. et al. Serine starvation induces stress and p53-dependent metabolic remodelling in cancer cells. Nature 493, 542–546, doi:10.1038/nature11743 (2013).

5 Maddocks, O. D. K. et al. Modulating the therapeutic response of tumours to dietary serine and glycine starvation. Nature 544, 372–376, doi:10.1038/nature22056 (2017).

6 Gravel, S. P. et al. Serine deprivation enhances antineoplastic activity of biguanides. Cancer Res 74, 7521–7533, doi:10.1158/0008-5472.CAN-14-2643-T (2014).

7 Kanarek, N. et al. Histidine catabolism is a major determinant of methotrexate sensitivity. Nature, doi:10.1038/s41586-018-0316-7 (2018).

8 Knott, S. R. V. et al. Asparagine bioavailability governs metastasis in a model of breast cancer. Nature 554, 378–381, doi:10.1038/nature25465 (2018).

9 Mentch, S. J. et al. Histone Methylation Dynamics and Gene Regulation Occur through the Sensing of One-Carbon Metabolism. Cell Metab 22, 861–873, doi:10.1016/j.cmet.2015.08.024 (2015).

10 Gao, X., Reid, M. A., Kong, M. & Locasale, J. W. Metabolic interactions with cancer epigenetics. Mol Aspects Med 54, 50–57, doi:10.1016/j.mam.2016.09.001 (2017).

11 Orentreich, N., Matias, J. R., DeFelice, A. & Zimmerman, J. A. Low methionine ingestion by rats extends life span. J Nutr 123, 269–274 (1993).

12 Lee, B. C. et al. Methionine restriction extends lifespan of Drosophila melanogaster under conditions of low amino-acid status. Nat Commun 5, 3592, doi:10.1038/ncomms4592 (2014).

13 Sun, L., Sadighi Akha, A. A., Miller, R. A. & Harper, J. M. Life-span extension in mice by preweaning food restriction and by methionine restriction in middle age. J Gerontol A Biol Sci Med Sci 64, 711–722, doi:10.1093/gerona/glp051 (2009).

14 Cabreiro, F. et al. Metformin retards aging in C. elegans by altering microbial folate and methionine metabolism. Cell 153, 228–239, doi:10.1016/j.cell.2013.02.035 (2013).

15 Miller, R. A. et al. Methionine-deficient diet extends mouse lifespan, slows immune and lens aging, alters glucose, T4, IGF-I and insulin levels, and increases hepatocyte MIF levels and stress resistance. Aging Cell 4, 119–125, doi:10.1111/j.1474-9726.2005.00152.x (2005).

16 Malloy, V. L. et al. Methionine restriction prevents the progression of hepatic steatosis in leptin-deficient obese mice. Metabolism 62, 1651–1661, doi:10.1016/j.metabol.2013.06.012 (2013).

17 Ables, G. P., Perrone, C. E., Orentreich, D. & Orentreich, N. Methionine-restricted C57BL/6J mice are resistant to diet-induced obesity and insulin resistance but have low bone density. PLoS One 7, e51357, doi:10.1371/journal.pone.0051357 (2012).

18 Ables, G. P. et al. Dietary methionine restriction in mice elicits an adaptive cardiovascular response to hyperhomocysteinemia. Sci Rep 5, 8886, doi:10.1038/srep08886 (2015).

19 Malloy, V. L. et al. Methionine restriction decreases visceral fat mass and preserves insulin action in aging male Fischer 344 rats independent of energy restriction. Aging Cell 5, 305–314, doi:10.1111/j.1474-9726.2006.00220.x (2006).

20 Ser, Z. et al. Targeting One Carbon Metabolism with an Antimetabolite Disrupts Pyrimidine Homeostasis and Induces Nucleotide Overflow. Cell Rep 15, 2367–2376, doi:10.1016/j.celrep.2016.05.035 (2016).

21 Miousse, I. R. et al. One-carbon metabolism and ionizing radiation: a multifaceted interaction. Biomol Concepts 8, 83–92, doi:10.1515/bmc-2017-0003 (2017).

22 Locasale, J. W. Serine, glycine and one-carbon units: cancer metabolism in full circle. Nat Rev Cancer 13, 572–583, doi:10.1038/nrc3557 (2013).

23 Hoffman, R. M. & Erbe, R. W. High in vivo rates of methionine biosynthesis in transformed human and malignant rat cells auxotrophic for methionine. Proc Natl Acad Sci U S A 73, 1523–1527 (1976).

24 Stern, P. H., Wallace, C. D. & Hoffman, R. M. Altered methionine metabolism occurs in all members of a set of diverse human tumor cell lines. J Cell Physiol 119, 29–34, doi:10.1002/jcp.1041190106 (1984).

25 Komninou, D., Leutzinger, Y., Reddy, B. S. & Richie, J. P., Jr. Methionine restriction inhibits colon carcinogenesis. Nutr Cancer 54, 202–208, doi:10.1207/s15327914nc5402_6 (2006).

26 Hens, J. R. et al. Methionine-restricted diet inhibits growth of MCF10AT1-derived mammary tumors by increasing cell cycle inhibitors in athymic nude mice. BMC Cancer 16, 349, doi:10.1186/s12885-016-2367-1 (2016).

27 Guo, H. et al. Therapeutic tumor-specific cell cycle block induced by methionine starvation in vivo. Cancer Res 53, 5676–5679 (1993).

28 Poirson-Bichat, F., Gonfalone, G., Bras-Goncalves, R. A., Dutrillaux, B. & Poupon, M. F. Growth of methionine-dependent human prostate cancer (PC-3) is inhibited by ethionine combined with methionine starvation. Br J Cancer 75, 1605–1612 (1997).

29 Sinha, R. et al. Dietary methionine restriction inhibits prostatic intraepithelial neoplasia in TRAMP mice. Prostate 74, 1663–1673, doi:10.1002/pros.22884 (2014).

30 Jeon, H. et al. Methionine deprivation suppresses triple-negative breast cancer metastasis in vitro and in vivo. Oncotarget 7, 67223–67234, doi:10.18632/oncotarget.11615 (2016).

31 Holter, N. S. et al. Fundamental patterns underlying gene expression profiles: simplicity from complexity. Proc Natl Acad Sci U S A 97, 8409–8414, doi:10.1073/pnas.150242097 (2000).

32 Byrne, A. T. et al. Interrogating open issues in cancer precision medicine with patient-derived xenografts. Nat Rev Cancer 17, 254–268, doi:10.1038/nrc.2016.140 (2017).

33 Hidalgo, M. et al. Patient-derived xenograft models: an emerging platform for translational cancer research. Cancer Discov 4, 998–1013, doi:10.1158/2159-8290.CD-14-0001 (2014).

34 Kim, M. K. et al. Characterization of an oxaliplatin sensitivity predictor in a preclinical murine model of colorectal cancer. Mol Cancer Ther 11, 1500–1509, doi:10.1158/1535-7163.MCT-11-0937 (2012).

35 Uronis, J. M. et al. Histological and molecular evaluation of patient-derived colorectal cancer explants. PLoS One 7, e38422, doi:10.1371/journal.pone.0038422 (2012).

36 Thiele, I. et al. A community-driven global reconstruction of human metabolism. Nat Biotechnol 31, 419–425, doi:10.1038/nbt.2488 (2013).

37 Saltz, L. B. et al. Bevacizumab in combination with oxaliplatin-based chemotherapy as first-line therapy in metastatic colorectal cancer: a randomized phase III study. J Clin Oncol 26, 2013–2019, doi:10.1200/JCO.2007.14.9930 (2008).

38 Douillard, J. Y. et al. Randomized, phase III trial of panitumumab with infusional fluorouracil, leucovorin, and oxaliplatin (FOLFOX4) versus FOLFOX4 alone as first-line treatment in patients with previously untreated metastatic colorectal cancer: the PRIME study. J Clin Oncol 28, 4697–4705, doi:10.1200/JCO.2009.27.4860 (2010).

39 Udofot, O. et al. Pharmacokinetic, biodistribution and therapeutic efficacy of 5-fluorouracil-loaded pH-sensitive PEGylated liposomal nanoparticles in HCT-116 tumor bearing mouse. J Nat Sci 2(2016).

40 Kirsch, D. G. et al. A spatially and temporally restricted mouse model of soft tissue sarcoma. Nat Med 13, 992–997, doi:10.1038/nm1602 (2007).

41 Sharpless, N. E. & Depinho, R. A. The mighty mouse: genetically engineered mouse models in cancer drug development. Nat Rev Drug Discov 5, 741–754, doi:10.1038/nrd2110 (2006).

42 Lee, C. L. et al. Generation of primary tumors with Flp recombinase in FRT-flanked p53 mice. Dis Model Mech 5, 397–402, doi:10.1242/dmm.009084 (2012).

43 Larrier, N. A., Czito, B. G. & Kirsch, D. G. Radiation Therapy for Soft Tissue Sarcoma: Indications and Controversies for Neoadjuvant Therapy, Adjuvant Therapy, Intraoperative Radiation Therapy, and Brachytherapy. Surg Oncol Clin N Am 25, 841–860, doi:10.1016/j.soc.2016.05.012 (2016).

44 Moding, E. J. et al. Atm deletion with dual recombinase technology preferentially radiosensitizes tumor endothelium. J Clin Invest 124, 3325–3338, doi:10.1172/JCI73932 (2014).

45 Moding, E. J. et al. Tumor cells, but not endothelial cells, mediate eradication of primary sarcomas by stereotactic body radiation therapy. Sci Transl Med 7, 278ra234, doi:10.1126/scitranslmed.aaa4214 (2015).

46 Durando, X. et al. Optimal methionine-free diet duration for nitrourea treatment: a Phase I clinical trial. Nutr Cancer 60, 23–30, doi:10.1080/01635580701525877 (2008).

47 Durando, X. et al. Dietary methionine restriction with FOLFOX regimen as first line therapy of metastatic colorectal cancer: a feasibility study. Oncology 78, 205–209, doi:10.1159/000313700 (2010).

48 Yuan, J., Bennett, B. D. & Rabinowitz, J. D. Kinetic flux profiling for quantitation of cellular metabolic fluxes. Nat Protoc 3, 1328–1340, doi:10.1038/nprot.2008.131 (2008).

